# The paradox of plant preference: the malaria vectors *Anopheles gambiae* and *Anopheles coluzzii* select suboptimal food sources for their survival and reproduction

**DOI:** 10.1101/2023.09.18.558223

**Authors:** Prisca S. L. Paré, Domonbabele F. D. S. Hien, Mariam Youba, Rakiswendé S. Yerbanga, Anna Cohuet, Louis-Clément Gouagna, Abdoulaye Diabaté, Rickard Ignell, Roch K. Dabiré, Olivier Gnankiné, Thierry Lefèvre

## Abstract

*Anopheles gambiae s.l.* mosquitoes, the primary malaria vectors in sub-Saharan Africa, exhibit selectivity among plant species as potential food sources. However, it remains unclear if their preference aligns with optimal nutrient intake and survival. Following an extensive screening of the effects of 31 plant species on *An. coluzzii* in Burkina Faso, we selected three species for their contrasting effects on mosquito survival, namely *Ixora coccinea, Caesalpinia pulcherrima and Combretum indicum*. We assessed the sugar content of these plants and their impact on mosquito fructose-positivity, survival, and insemination rate, using *Anopheles coluzzii* and *Anopheles gambiae*, and with glucose 5% and water as controls. Plants displayed varying sugar content and differentially affected the survival, sugar intake and insemination rate of mosquitoes. All three plants were more attractive to mosquitoes than controls, with *An. gambiae* being more responsive than *An. coluzzii*. Notably, *C. indicum* was the most attractive but had the lowest sugar content and offered the lowest survival, insemination rate, and fructose positivity. Our findings unveil a performance-preference mismatch in *An. coluzzii* and *An. gambiae* regarding plant food sources. Several possible reasons for this negative correlation between performance and preference are discussed.

## Introduction

According to optimal foraging theory, animals are expected to be able to discriminate and select among food resources that maximize nutrient intake and overall fitness, while considering the physiological and ecological trade-offs associated with foraging (*e.g*., time, exposure to enemies), as well as ingestion and processing of food ^1,2^. In choice situations, numerous organisms - ranging from slime molds to humans - are capable of selecting food resources that provide the optimal balance of energy and nutrients, resulting in improved fitness ^3–10^. As important pollinators, biocontrol agents, or pests, sugar-feeding insects, such as bees, butterflies, parasitoid wasps, or adult flies have received significant attention for their ability to self-select food resources that enhance their survival and reproductive success ^11–19^. Dipteran insects, including mosquitoes, black flies, sand flies, and biting midges, can transmit pathogens of major public and animal health concern. These vectors often visit flowers to gather nectar as a source of energy. However, it remains unclear whether the preference of insect vectors among different nectar sources correlates positively with the fitness benefits offered by the selected plants.

Mosquitoes are well-known for feeding on blood, during which they can transmit pathogens responsible for devastating diseases, such as malaria, dengue fever or Zika ^20^. In most species, the females are hematophagous and blood is an essential source of protein to produce their eggs. A growing body of research, however, shows that, just like males, plant fluids, including floral and extra-floral nectar, fruit, honeydew and phloem or xylem sap, are an essential resource for mosquito females, with critical epidemiological implications ^21^. As the primary energy source for many important mosquito vectors, including *Culex*, *Aedes*, and *Anopheles*, plant fluids not only fuels flight and activity, but can also be an important determinant of longevity and reproductive success (reviewed in ^22^). Previous studies have shown that changes in the abundance or composition of particular flowering plant species can exert significant effects on the population dynamic and vectorial capacity of *Anopheles gambiae sensu lato*, the primary vector of the malaria parasite, *Plasmodium falciparum*, in Africa, through their impacts on the lifespan, susceptibility to pathogens, and reproductive output of individual mosquitoes ^23–25^.

Given the importance of floral resources for the biology of major malaria vectors, we still know surprisingly little about how intra- and inter-specific variations in plant quality affects mosquito performance. Nectar, the main food resource that mosquitoes exploit from plants, is mostly composed of sugars (sucrose and its monomers, glucose and fructose) and, to a lesser extent, primary metabolites as amino acids, lipids, vitamins, proteins, and secondary metabolites, such as terpenes, alkaloids and phenolics ^26,27^. Nectar quality varies among flowering plant species with respect to the concentration and composition of these sugars and other constituents ^28,29^. Therefore, not all flowering plants are expected to offer suitable resources in terms of energy and nutrient intake and hence of fitness benefits to malaria vectors, which, in turn, should likely lead to some degree of variability in their attractiveness to mosquitoes. Although several studies have examined flower suitability by measuring the fitness components of individual mosquitoes provided with a variety of plant species, only few have attempted to establish a connection with mosquito foraging preferences ^22^.

Mosquitoes rely on the integration of olfactory, visual, and gustatory cues to locate and select their host plants ^22,29^. Based on observations gathered from field, semi-field, and laboratory settings, it appears that, although malaria vectors can use a wide variety of plant species as food sources, they display some degree of selectivity among the different plant species available in their natural habitats ^30–36^. To the best of our knowledge, only three studies have investigated whether *An. gambiae s.l.* exhibit a significant preference for the plant species that best support nutrient intake, survival, and/or fecundity (*i.e*., positive performance-preference relationships). First, among three flowering plant species in La Reunion Island, male *Anopheles arabiensis* were observed to accumulate greater levels of energy reserves, including sugar, lipids, and proteins, when feeding on their most preferred plant species ^31^. Second, following the classification of 13 plant species according to their attractiveness to female *An. gambiae* in Kenya ^32^, mosquitoes were fed with one of each of five of the most attractive plants and one of the least attractive ^37^. Although four of the five most attractive plants provided greater female survival and fecundity compared to the least attractive species, the highly attractive *Parthenium hysterophorus* provided low survival and fecundity ^37^. Thirdly, among four plant species that were highly attractive to *An. gambiae* females ^34^, three exhibited high sugar content and led to greater survival rates, while *P. hysterophorus* acted as a deceptive trap, causing high mortality probably associated with a low sugar content ^38^. Collectively, these results suggest that, with the exception of *P. hysterophorus*, mosquitoes obtain greater fitness benefits when feeding on their preferred plant species, presumably due to the high sugar content present in these preferred plants. While the previous body of research was carried out in La Reunion and Kenya using *An. gambiae* and *An. arabiensis*, comparable studies in other parts of Africa and on other important vectors, such as *An. coluzzii*, are lacking. Expanding our understanding of the nutritional ecology of the major malaria vectors in Africa will provide insights into the factors that shape vectorial capacity in different endemic settings. This may also contribute to the identification of new attractive odor blends that could be used, for example, in the development of sugar baits/traps.

To gain further perspectives on the nature of malaria vector-plant interactions, we tested performance-preference relationships in a series of laboratory experiments in Burkina Faso using both *An. gambiae* and *An. coluzzii*, two primary vectors of *P. falciparum* in West Africa. We hypothesized that mosquito females would exhibit a significant selective behavior for the plant species that best support their survival and that feeding preference could be mediated by sugar intake and sugar content in plants. First, we screened the effects of 31 flowering plant species on *An. coluzzii* and selected three species with contrasting effects on mosquito survival. Second, we used no-choice feeding assays to assess whether the consumption of these three plants affected the survival and sugar intake of *An. gambiae* and *An. coluzzii* females, using water and a 5% glucose solution as controls. Third, we quantified the sugar content of these three plant species. Owing to its effect on the body condition of both males and females, the variability in sugar quality among plant species can cause a proportion of females to remain uninseminated and therefore reduce the egg output of a population. Fourth, we therefore assessed the effect of the consumption of each of these three plant species, either by males alone, by females alone or by both sexes on the insemination rate. Fifth, we performed multiple-choice behavioral assays to determine whether females preferentially select plant species that offer higher fitness benefits in terms of sugar intake, insemination rates and/or survival.

## Methods

### Mosquito strains

Laboratory-reared *An. coluzzii* and *An. gambiae* were obtained from outbred colonies established in 2019, that have since been repeatedly replenished with wild-caught gravid females collected in the Vallée du Kou (11°23’N, 4°24’W) and Soumousso (11°04’N, 4°03’W), respectively, in south-western Burkina Faso, and identified by SINE PCR ^39^. Females were maintained on rabbit blood by direct feeding (protocol approved by the national committee of Burkina Faso; IRB registration #00004738 and FWA 00007038) for egg production. Larvae were reared in 1 l of tap water in plastic trays and fed daily with TetraMin® Baby Fish Food (Tetrawerke, Melle, Germany) until adulthood. The adult mosquitoes were held in 30 cm × 30 cm × 30 cm mesh-covered cages under standard controlled conditions (27 ± 2 °C, 70 ± 5% RH and at a 12 h: 12 h light: dark rhythm), and emerged males and females were fed daily with 5% glucose.

#### 1. Inter-specific plant effects on mosquito performance

##### Experiment 1.1: Screening of plant species that differentially support mosquito survival

The effect of 31 flowering plant species (Supp. Fig. 1) on the survival of *An. coluzzii* mosquitoes was evaluated. The plants were selected based on their presence around human dwellings and public areas of the city of Bobo-Dioulasso and the village of Farako-ba, two localities in western Burkina Faso. The flowers were collected daily between 3 pm and 4 pm and offered to mosquitoes in 30 cm × 30 cm × 30 cm mesh-covered cages at 5 pm. Between seven and ten freshly cut stems of flowering plants were arranged in a bouquet (with leaves removed) and introduced into the cages. The base of the bouquet was wrapped in moistened paper towels and covered with an aluminum sheet so that mosquitoes had no access to the moistened paper ^25,40^. A 5% glucose solution was used as a positive control by soaking a cotton pad with this solution and placing it on top of the control cage. The flower bouquets and the 5% glucose cotton pad were changed daily at 5 pm. Disposable plastic cups (20 cl) containing about 30-40 pupae (males and females) were randomly assigned to one plant species. Adult mosquitoes (males and females with equal sex-ratio, exact sample sizes are indicated in the legend of the corresponding figure below) emerging from these pupae were kept for 6 consecutive days on their assigned diet (one of four flowers or the glucose control solution). Mosquito survival was recorded from day 1 to day 6 post-emergence. This consisted in counting dead mosquitoes daily, regardless of sex, between 4 and 5 pm, and removing them from the cages. A specific permit for the sampling of the 31 plant species was obtained from the Ministère de l’Environnement, de l’Energie, de l’Eau et de l’Assainissement des Hauts-Bassins of Burkina Faso. All plant species were identified by a botanist from the “Institut de Recherche pour le Développement’’ followed by an independent confirmation by a phytoecologist from the University Joseph KI-ZERBO/University Center of Ziniaré Burkina Faso according to a catalogue done by Thiombiano et al. ^41^.

To further investigate the influence of plant species on mosquito survival rate, insemination rate and sugar positivity, two experiments were conducted.

##### Experiment 1.2: survival, cold-anthrone tests, determination of the degree Brix and the total sugar content

Based on the screening experiment 1.1, three plant species were selected for their contrasting effects on mosquito survival (ranging from positive to negative effects on survival, Fig. 1, Supp. Table 1), *Ixora coccinea* L. (Rubiaceae)*, Caesalpinia pulcherrima* (L.) Sw. (Fabaceae-Caesalpinioideae), and *Combretum indicum* (L.) Jongkind (Combretaceae*). Ixora coccinea* and (i) *C. pulcherrima* provided high and medium mosquito survival, respectively, whereas *C. indicum* induced poor survival. The other species provided equivalent survival to *I. coccinea* but were not selected for the subsequent experiments because they were less abundant and available than *I. coccinea* at the time of the experiments. This was also true for the other species that provided equally poor survival as *C. indicum.* Specimens of *C. pulcherrima*, *C. indicum* and *I. coccinea*, were deposited in the herbarium of the Nazi Boni University, Bobo-Dioulasso, Burkina Faso, under the identification numbers UNB-947, UNB-948, and UNB-949, respectively. All three species are exotic, widely cultivated ornamental plants in cities of Burkina Faso. *Ixora coccinea,* the jungle geranium, and *C. indicium*, the rangoon creeper, are native to tropical Asia. *Caesalpinia pulcherrima,* the peacock flower, has a (sub-) tropical distribution but its exact origin remains unclear ^42^.

**Fig. 1:**
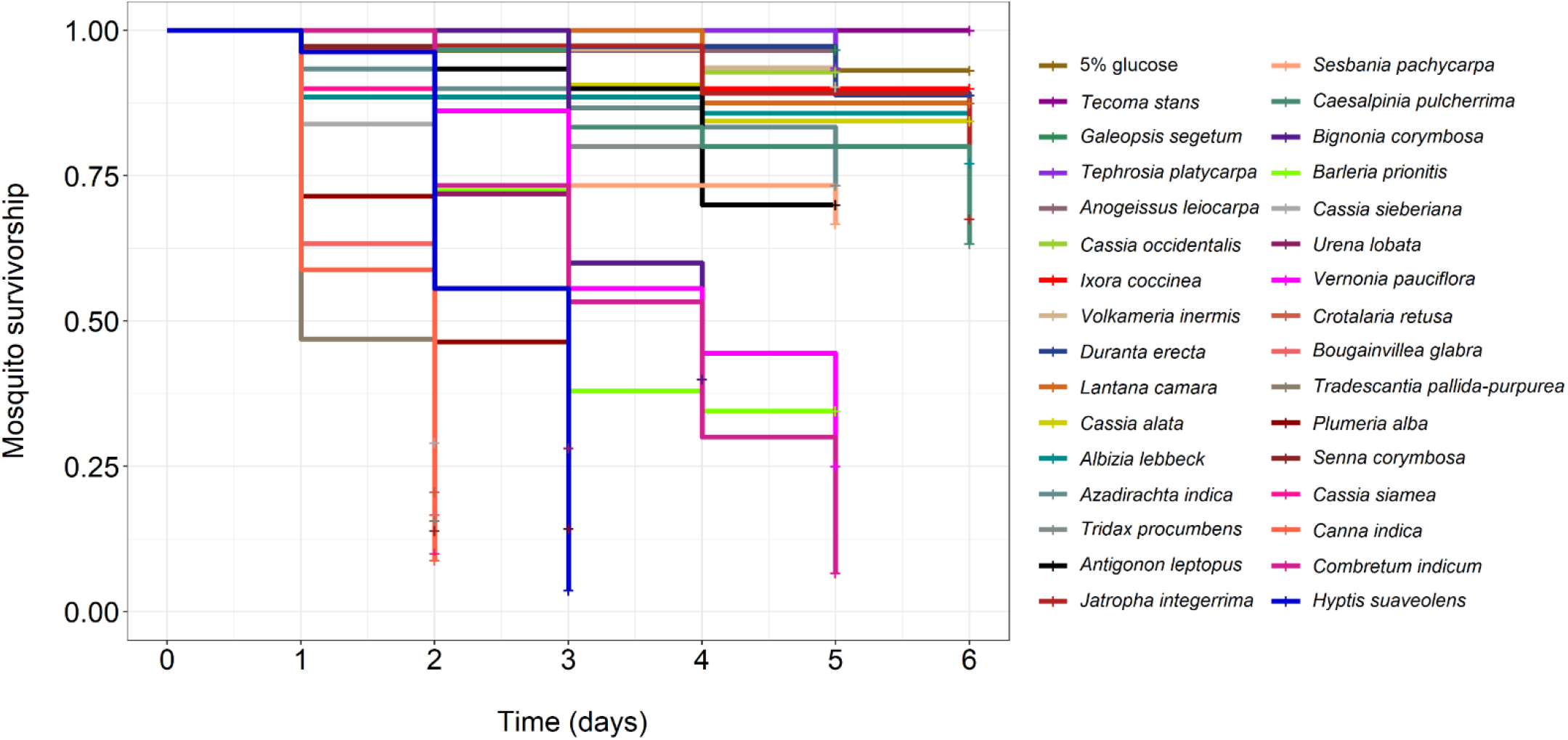
The effect of 31 plant species and a 5% control glucose solution on the survival of *An. coluzzii*. Kaplan–Meier curves represent the proportion of live mosquitoes over time for each treatment. Between 27 and 37 emerging mosquitoes (males and females, mean ± se: 31.25 ± 0.45, median: 30) were maintained on one of the treatment and survival was recorded from day 1 to day 6 post-emergence.

To further investigate the effect of these three plant species on the survival of malaria vectors, unlike the screening experiment 1.1, both *An. coluzzii* and *An. gambiae* were used, and males and females were distinguished in this experiment. A total of 20 cages were used for this test (2 cages per treatment and species) each containing 130 nymphs of both sexes. Upon emergence, males and females of each mosquito species were maintained together on one of five treatments: *I. coccinea*, *C. pulcherrima*, *C. indicum*, water (negative control) and 5% glucose solution (positive control) (Supp. Fig. 2). Mosquitoes were exposed to these treatments in the same manner and timing as in the screening experiment 1.1. The mortality of males and females was monitored every day between 4 pm and 5 pm until all mosquitoes were dead.

In parallel with the longevity experiment, 10 additional 20 cm x 20 cm x 20 cm cages (1 cage per treatment and species) were set up to perform cold-anthrone tests ^43^ (Supp. Fig. 3). One hundred and ten nymphs were introduced into each cage at 9 am. The following day after emergence, at 17 pm, the mosquitoes were provided with one of the five treatments (water, 5% glucose, *I. coccinea, C. pulcherrima, C. indicum*) as described above. On the second day after emergence, at 8 am (i.e., after 15 hours of exposure to the treatments), all mosquitoes were aspirated to determine the proportion of fructose-positive mosquitoes.

The °Bx and the total sugar content of the flowers of *I. coccinea, C. pulcherrima* and *C. indicum* were determined in the biochemistry and microbiology laboratory of the Département Technologie Alimentaire of the Institut de Recherche en Sciences Appliquées et Technologies of Bobo-Dioulasso. A hand-held refractometer (Atago, ATC, Tokyo, Japan) with a scale of 0-32%, and an accuracy of 0.2%, was used for the determination of °Bx ^44^. Briefly, the flowers of each plant species were crushed using an electric grinder, type A11 basic (IKA^®^-Werke GmbH & Co. KG, Staufen, Germany). One gram of crushed material of each species was wrapped in a net and squeezed to obtain a liquid solution. The refractometer was calibrated by placing 2 drops of distilled water on the main prism and was then cleaned and dried after each calibration. The solutions obtained by grinding were then placed on the refractometer prism. For each plant species, three measurements were made, each on different dates, and using different plant individuals.

Total sugar content was determined according to the sulfuric orcinol colorimetric method^45^. In the presence of concentrated sulfuric acid and at high temperature, the hexoses and pentoses of the medium undergo a thorough internal dehydration, followed by a cyclization leading to the formation of furfural and 5-hydroxymethylfurfural derivatives, reacting with orcinol to give a yellow-red complex. This allows the total sugar concentration of the sample to be monitored by reading the absorbance at 510 nm. At 6 am, flowers of each plant species were collected and brought back to the laboratory where they were immediately crushed by species using an electric grinder. One g of crushed material from each species was taken and introduced into a 100 ml volumetric flask to which 5 ml of distilled water was added. The mixture was put under magnetic stirring for 10 min and the volume was completed to 100 ml with distilled water. One ml of each solution was introduced into test tubes (*i.e.*, two tubes for each individual plant species) to which 2 ml of sulfuric orcinol reagent and 7 ml of 60% H_2_SO_4_ solution were added. The mixture was incubated in a boiling water bath for 20 min, placed under ambient temperature in the dark for 45 min and then in natural light for 10 min. Then, a dilution was performed by adding 2 ml of 60% H_2_SO_4_ in 1 ml of each sampled solution. The optical densities of each diluted solution were read at 510 nm using a spectrophotometer (V-1100D spectrophotometer, J.P.SELECTA, Barcelona, Spain). Sugar concentrations were determined using a standard curve, with D-glucose as the reference sugar. The total sugar content of each species was expressed in D-glucose equivalent as g/100 g fresh material ^45^.

##### Experiment 1.3: Survival and insemination

To evaluate the effect of plant diet on mosquito survival and insemination rates, three assays, each with a different design (Supp. Fig. 4), were conducted in parallel using *An. coluzzii* and *An. gambiae*. The three assays were replicated three times (*i.e*., a total of three experimental replicates were performed). The objective of design 1 was to determine whether the treatments, provided to both males and females, can influence sexual performance, as measured by female insemination rate. Design 2 aimed to assess the impact that the plants might have on the ability of females to get inseminated by males. In contrast, design 3 tested the effect of treatments on the ability of males to inseminate females.

- Design 1: Circa 30 newly emerged males and 30 newly emerged females were introduced into 20 cm × 20 cm × 20 cm cages and kept together for 5 days on one of four treatments: *I. coccinea, C. pulcherrima, C. indicum* and 5% glucose solution (Supp. Fig. 4a). Sample sizes varied slightly across species and replicates according to mosquito availability in the insectary (the exact sample sizes for each treatment, species and replicates are indicated in Supp. Table 2).
- Design 2: Circa 30 newly emerged females were kept in 20 cm × 20 cm × 20 cm cages and fed daily with one of the four treatments. Three days later at 8 am, circa 30 males of the same age as the females, and previously fed on a 5% glucose solution for 3 days, were introduced into the female cages. Males and females were then kept together for 2 days on their assigned treatment (Supp. Fig. 4b).
- Design 3: This design was similar to design 2 except that circa 30 newly emerged males, instead of females, were fed daily with one of the four treatments. On the morning of the 3rd day at 8 am, circa 30 females, previously fed on a 5% glucose solution for 3 days, were introduced into the male cages. Males and females were kept together for 2 days on their assigned treatment (Supp. Fig. 4c).

Of particular note, males and females were housed together for 5 days in design 1, while in designs 2 and 3 the duration of contact between males and females was 3 days only. For each assay, plants were replaced every day with fresh materials in the same manner and timing as in the previous experiments. On the fifth day of the experiment at 8 am, the remaining females were retrieved and anesthetized at −20°C for three minutes. Spermathecae were dissected under a stereomicroscope (LEICA^®^ S9E, Wetzlar, Germany) in a drop of distilled water and mounted under a coverslip. A gentle pressure was exerted on the coverslip with dissecting forceps to rupture the spermatheca, which was then observed under a compound light microscope (LEICA DM1000 LED, Germany) at 400× magnification to assess the insemination status (Supp. Fig. 5). In all assays, mosquito mortality was recorded every day at 8 am until day 5 and all remaining live male and female mosquitoes (used for spermatheca dissection) were considered in the survival analysis and given a censoring indicator of “0”. However, when either sex was first maintained on 5% glucose for 3 days (*i.e*., males in design 2 and females in design 3), then survival was monitored only from day 3 post-emergence.

#### 2. Mosquito behavioral response to plant species

Behavioral analysis was conducted in a multiple-choice experimental device consisting of four large insect release cages (1 m × 1 m × 1 m) set-up in a climate controlled room (29 °C, 70 ± 5% RH). Each large release cage contained five smaller cages of 15 cm × 15 cm × 15 cm, each housing one of the five treatments (traps) (Supp. Fig. 6a, b). The mesh screen of these traps was raised 3 cm from the ground to allow mosquitoes to enter (Supp. Fig. 6c).

On the day of the behavioral test at 8 am, between 16-108 females of *An. coluzzii* (mean: 59.38 ± 1.20, median: 60.50) and between 35-109 females of *An. gambiae* (mean: 71.50 ± 0.91, median 70.00), aged between one and four days, and previously maintained on 5% glucose, were aspirated and introduced into cardboard cups (four cups per species). The number of mosquitoes introduced into the cups (sample sizes) varied across behavioral tests (replicates) depending on mosquito availability in the insectary. To examine possible differences in behavioral responses between *An. gambiae* and *An. coluzzii*, two cups (one of each species) were released simultaneously into each large release cage. Females of both species were therefore exposed to the same flower bouquets, hence preventing the possible confounding effect of mosquito species and individual plant factors. For this purpose, one of the two mosquito species was marked with colored powder (Luminous Powder Kit, Bioquip Products Inc 2321 Gladwick Street Rancho Dominguez, CA 90220, USA). Marking was alternated between species, cages and replicates to avoid confounding factors. The release of one cup of unmarked *An. gambiae* and one cup of marked *An. coluzzii* simultaneously in the same cage (and vice versa in other cages) allowed the species to be distinguished. Because mosquito number (varying sample sizes) and color marking might influence behavioral response, density and color were considered in the statistical analyses. A cotton pad soaked with water was placed on the mosquito cups, which were then kept under insectary conditions (27 ± 2°C and 70 ± 5 % RH) prior the test.

At 3.30 pm on the day of the behavioral test, flowers of *I. coccinea, C. pulcherrima, C. indicum* were collected and made into a flower bouquet as described above, and then positioned in the small trap cages. In addition, two traps were baited with either 5% glucose cotton pads or water pads. These cotton pads were placed on a 20 cl disposable plastic cup positioned upside down on the bottom of the trap cages. A total of 20 small trap cages were used during each behavioral test (5 traps × 4 large cages). Each of these traps contained either the flower bouquet of *I. coccinea, C. pulcherrima or C. indicum,* or the 20 cl disposable cup holding either the 5% glucose cotton pad or the water cotton pad (Supp. Fig. 7). The position of the traps was alternated randomly within the cages and among the 16 releasing nights (replicates).

At 6 pm, *An. gambiae* and *An. coluzzii* contained in either of two cups (starved of sugar for a period of 10h) were released simultaneously into one of the four release cages, and were allowed 12 h to make a choice between treatments. Mosquitoes that were attracted to a treatment entered the cage trap through the 3 cm gap. With this configuration, the possibility that mosquitoes that entered a given trap managed to exit and remained in the release cage or visited another trap cannot be excluded. The following day the mosquitoes were collected. First, the odor trap nets were gently lowered and then tied to prevent the trapped mosquitoes from escaping. Mosquitoes that did not make a choice, *i.e*., those remaining in the large release cages, were aspirated and stored in cardboard cups. Then, the nets of the large cages were removed and the caught mosquitoes were aspired from the small trap cages. These mosquitoes were placed in disposable plastic cups corresponding to their treatment and release cage identity. All aspirated mosquitoes were anaesthetized at −20°C for counting by species (based on the color), by large release cage and by trap cage (treatment). The 12h duration of the behavioral trial was chosen on the basis of previous studies, which revealed that sugar feeding in both male and female *An. gambiae* followed a largely unimodal crepuscular/nocturnal diel rhythm ^46^.

The behavioral assay was repeated 16 times (16 nights) for a total of 64 choice episodes (16 nights × 4 large cages) for each mosquito species. Two traits were measured to analyse mosquito behavioral responses:

i. activation rate, which is the number of mosquitoes caught in all traps out of the total number of mosquitoes released and,
ii. plant relative attractiveness, which is the number of mosquitoes caught in each odor trap out of the total number of mosquitoes caught in all odor traps.

### Statistical analysis

All statistical analyses were performed using R software (version 4.0.5) ^47^. Cox proportional hazard model (“coxph” function of the “survival” library version 3.2-10 ^48^) with censoring was performed to test the effect of diet (32 levels) on mosquito survivorship (Experiment 1.1). A Cox’s proportional hazard mixed regression model (“coxme” library version 2.2-16 ^49^) without censoring and with cage (20 levels) set as a random effect was performed to test the effect of treatment (5 levels: water, 5% glucose, *I. coccinea, C. pulcherrima, C. indicum*), mosquito species (2 levels: *An. coluzzii* and *An. gambiae*), sex (2 levels) and their interactions on mosquito survivorship (Experiment 1.2). Logistic regression by generalized linear model (GLM, quasibinomial errors, logit link) was used to test the effect of treatment, mosquito species, sex, and their interactions on the proportion of mosquitoes positive to cold anthrone (Experiment 1.2). The effect of treatment (4 levels: 5% glucose, *I. coccinea, C. pulcherrima, C. indicum*), mosquito species (2 levels), design (3 levels), sex (2 levels) and their interactions on mosquito survivorship was analysed using a censored Cox’s proportional hazard mixed regression model (“coxme” library version 2.2-16 ^49^) with replicate (3 levels) set as a random effect (Experiment 1.3). The effect of treatment, mosquito species, design, and their interactions on insemination rate was analysed using a logistic regression by generalized mixed linear models (GLMM, binomial errors, logit link; “lme4” library version 1.1-32 ^50^) with replicate (3 levels) set as a random effect (Experiment 1.3). A binomial GLMM was also used to test the effect of species (2 levels), coloration (2 levels: uncolored and colored), density (the number of mosquitoes released in the large cages, log-transformed) and relevant two-ways interactions on mosquito activation (Experiment 2). A mixed-effects multinomial logistic regression model (“mblogit” function of the “mclogit” library version 0.9.6 ^51^) was used to explore the effect of species (2 levels), coloration (2 levels), density (log-transformed), and relevant two-ways interactions on plant relative attractiveness to mosquitoes (Experiment 2). The relative odds ratios were then derived to compare the likelihood of mosquitoes choosing one treatment over another. In these two mixed models (binomial GLMM for mosquito activation and multinomial GLMM for plant relative attractiveness), the cage (4 levels) was nested within night (16 levels) and considered together as nested random effects.

For all analyses, the “Anova” function from the “car” library version 3.1-1 ^52^ was used to estimate the significance of terms except for the attractiveness analysis (multinomial GLMM using mclogit) for which the best model was selected based on the Akaike Information Criteria (AIC). Multiple pair-wise post-hoc tests were performed to compare each level of the treatment when the latter was significant using the “emmeans” function of the “emmeans” library version 1.5.5-1 ^53^.

## Results

### 1. Inter-specific plant effects on mosquito performance

#### Experiment 1.1: Screening of plant species that differentially support mosquito survival

Mosquito survival rate varied widely among plant species (LRT *X^2^*_31_ = 869.14; P < 0.001, Fig. 1, Supp. Table 1). On day 6 post-emergence, when mortality monitoring was stopped, the survival rate of mosquitoes fed with the 5% control glucose solution was 93 ± 0.1%. On the basis of multiple pairwise post-hoc comparisons, the survival rate of mosquitoes fed with either *Tecoma stans* (100 ± 0.00%), *Galeopis segetum* (97 ± 0.07%)*, Tephrosia platycarpa* (94 ± 0.09%), *Cassia occidentalis* (93 ± 0.1%), *Anogeissus leiocarpa* (93 ± 0.1%), *Volkameria inermis* (90 ± 0.11%), *I. coccinea* (90 ± 0.11%), *Duranta erecta* (89 ± 0.11%), *Lantana camara* (88 ± 0.12%), *Cassia alata* (84 ± 0.14%), *Albizia lebbeck* (77 ± 0.16%), *Azadirachta indica* (73 ± 0.18%), *Tridax procumbens* (73 ± 0.18%), *Antigonon leptopus* (70 ± 0.20%), *Jatropha integerrima* (68 ± 0.18%), *Sesbania pachycarpa* (67 ± 0.21%) or *C. pulcherrima* (63 ± 0.22%) was similar to that of mosquitoes fed with the 5% glucose solution (Supp. Table 3). In contrast, plant species, including *Bignonia corymbosa* (40 ± 28%), *Barleria prionitis* (34 ± 29%), *Cassia sieberiana* (29 ± 30%), *Urena lobata* (28 ± 29%), *Vernonia pauciflora* (25 ± 28%), *Crotalaria retusa* (21 ± 30%), *Bougainvillea glabra* (17 ± 33%), *Tradescantia pallida-purpurea* (16 ± 32%), *Plumeria alba* (14 ± 34%), *Senna corymbosa* (14 ± 30%), *Cassia siamea* (10 ± 34%), *Canna indica* (9 ± 32%), *C. indicum* (7 ± 35%) and *Hyptis suaveolens* (4 ± 37%) negatively impacted mosquito survival compared to the 5% glucose solution (Fig. 1, Supp. Table 3).

To assess performance-preference relationships, we subsequently selected three species that were abundant at the time of the following experiments, and had contrasting effects on mosquito survival, namely *I. coccinea*, *C. pulcherrima* and *C. indicum* (ranked according to their effect on survival from positive to negative).

#### Experiment 1.2: Survival, cold-anthrone tests, determination of the degree Brix and total sugar content

##### Survival

While there was no species effect on mosquito survival (LRT *X^2^*_1_ = 2.5, P = 0.11, Fig. 2, Supp. Tables 4, 5), females exhibited a median survival time of 10 days, 4 days greater than that of males (LRT *X^2^*_1_ = 10.6, P = 0.001, Fig. 2). Mosquito survival strongly varied among treatments (LRT *X^2^*_4_ = 861, P < 0.001, Fig. 2, Supp. Tables 4, 5), and all pairwise differences were significant except that between *I. coccinea* and *C. pulcherrima* (Supp. Table 6). In the absence of a food source, *i.e.,* water only, mosquitoes died within 3-5 days and were 135.65 times more likely to survive when kept on a 5% glucose solution rather than on water (Fig. 2, Table 1). Compared to the water control, the chances of survival were 20.69, 38.19 and 42.60 times greater when mosquitoes were maintained on *C. indicum, C. pulcherrima* and *I. coccinea*, respectively (Table 1). There was an interaction between treatment and species (LRT *X^2^*_4_ = 35, P < 0.001, Fig. 2), with a survival difference between *C. indicum* and *C. pulcherrima* within *An. gambiae* but not within *An. coluzzii*. There also was a treatment by sex (LRT *X^2^*_4_ = 21, P = 0.003) and a species by sex (LRT *X^2^*_1_ = 4, P = 0.04) interaction. There was no three-way interaction (Fig. 2, Supp. Table 4).

**Fig. 2:**
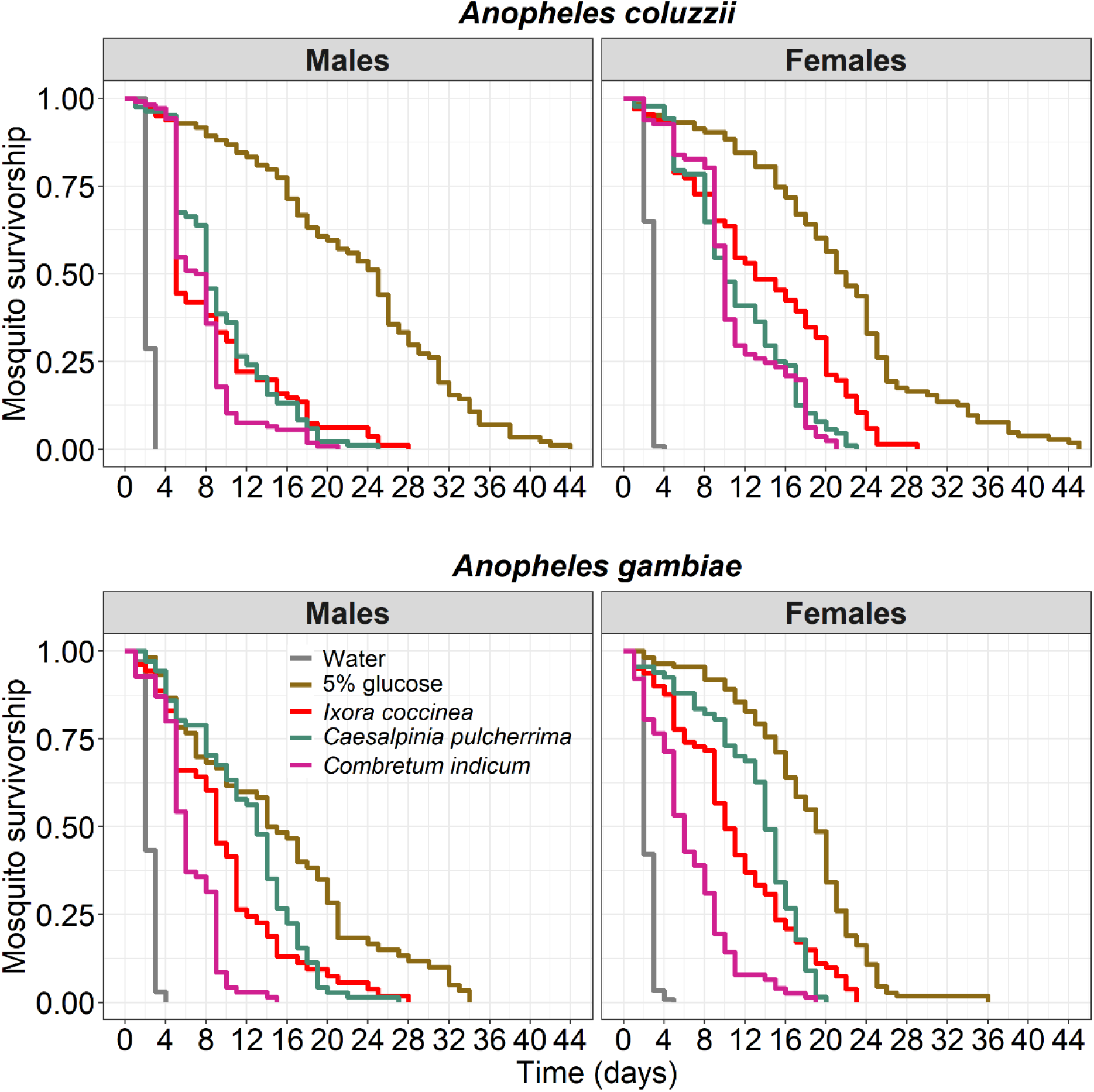
Effect of diet on the survival of *Anopheles coluzzii* and *Anopheles gambiae* according to sex. Kaplan–Meier curves represent the proportion of live mosquitoes over time for each diet (water = negative control, 5% glucose solution = positive control). There were 130 nymphs in each cage. The survival of males and females *An. coluzzii* and *An. gambiae* were monitored until all mosquitoes were dead.

**Table 1:** Risk of mosquito survival (hazard ratio) without censoring for each treatment relative to the control (water). lower .95 - upper .95 represents the 95% confidence interval around the hazard ratio.

| Diet | Hazard ratio (lower .95 - upper .95) | Z | P-value |
| --- | --- | --- | --- |
| 5% glucose | 135.65 (0.01-0.01) | 35.41 | < 0.001 |
| *Ixora coccinea* | 42.60 (0.02-0.03) | 28.35 | < 0.001 |
| *Caesalpinia pulcherrima* | 38.19 (0.02-0.03) | 28.01 | < 0.001 |
| *Combretum indicum* | 20.69 (0.04-0.06) | 24.31 | < 0.001 |

##### Cold-anthrone test

The anthrone test was carried out on all mosquitoes that emerged after 15 hours of contact with the treatments. Overall, the proportion of fructose-positive mosquitoes varied among treatments (LRT *X^2^*_4_ = 267.24, P < 0.001, Fig. 3, Supp. Table 7), ranging from 0% in water and in glucose, to 6 ± 0.15% in *C. indicum*, 13 ± 0.13% in *C. pulcherrima*, and 50 ± 0.10% in *I. coccinea*. Fructose positivity did not vary between species and between sex (12 ± 0.08% and 16 ± 0.09% for *An. coluzzii* and *An. gambiae*, respectively, LRT *X^2^*_1_ = 2.78, P = 0.1; 11 ± 0.09%, and 16 ± 0.08% for males and females, respectively, LRT *X^2^*_1_ = 2.96, P = 0.09, Fig. 3, Supp. Table 7). There was a significant interaction between treatment and species (LRT *X^2^*_4_ = 12.66, P = 0.01, Fig. 3), with a higher proportion of fructose-positive individuals in the *C. pulcherrima* treatment compared to *C. indicum* for *An. gambiae*, while no such difference was noted for *An. coluzzii*.

**Fig. 3:**
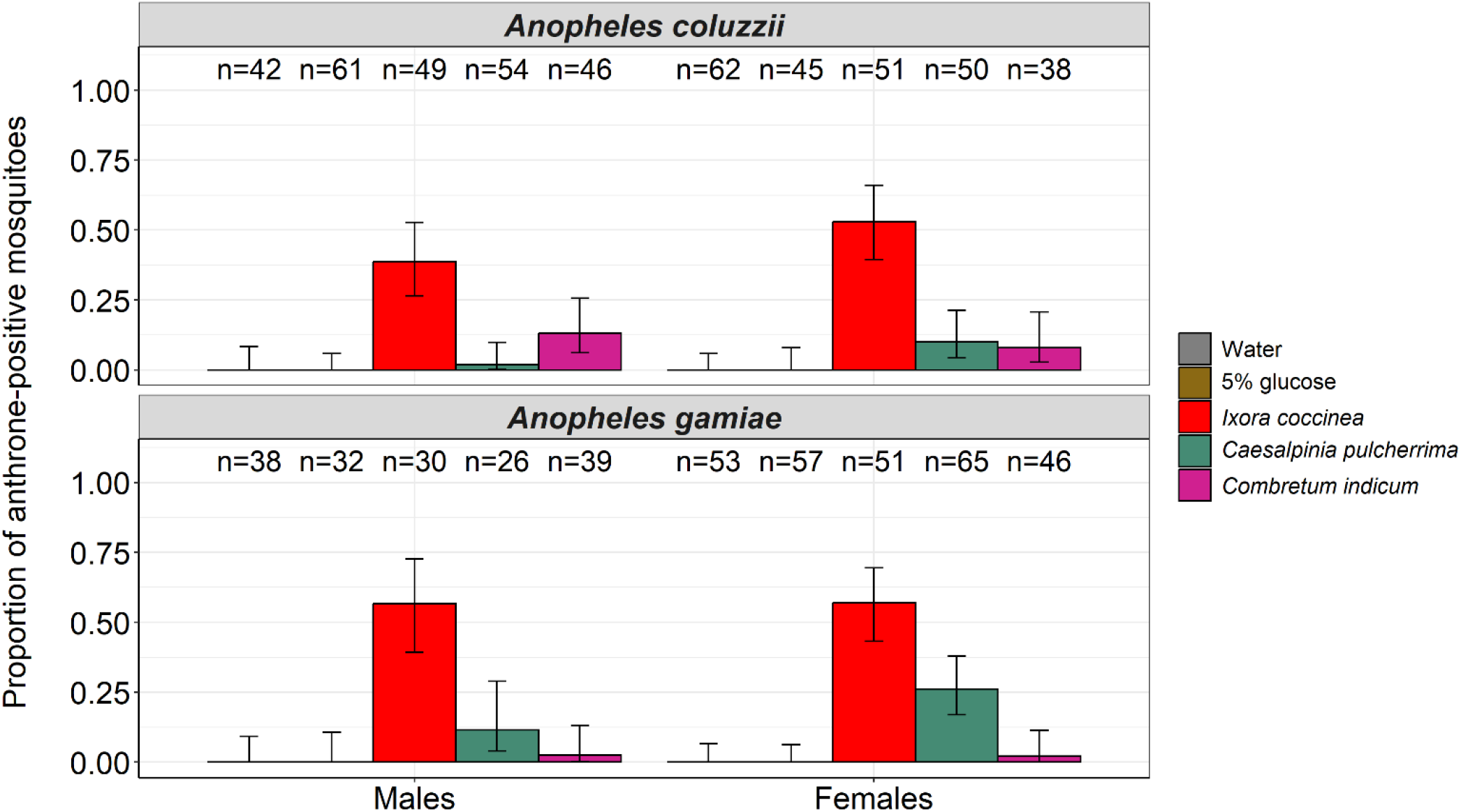
Proportion of *Anopheles coluzzii* and *Anopheles gambiae* (males and females) tested positive to fructose after exposure to one of five treatments: water, 5% glucose solution, *I. coccinea, C. pulcherrima and C. indicum* for 15 h for *An. coluzzii* and *An. gambiae*. The numbers above the barplots represent the sample size for each treatment. The error bars represent the variability of data with 95% confidence interval (±95% CI).

##### Determination of the degree Brix and the total sugars

Although there were only 3 observations per plant species, thus precluding statistical analysis, our results suggest that the °Bx of *C. pulcherrima* was higher than that of *C. indicum* and *I. coccinea* (Table 2). With respect to sugar content, only two observations per plant were available, but *C. indicum* demonstrated values half of that of *C. pulcherrima* and *I. coccinea* (Table 2).

**Table 2:** Mean °Bx and sugar content. The mean °Bx value was calculated based on three measurements taken from three different flower/plant individuals at different time intervals, whereas the mean sugar content was determined by averaging two measurements taken from the same plant during the same time period. Sugar content was expressed in g/100 g of fresh matter. se: standard error.

| Plants | Mean °Bx (±se) | Mean sugar content (±se) |
| --- | --- | --- |
| *Ixora coccinea* | 12.67 ± 0.77 | 8.06 ± 0.68 |
| *Caesalpinia pulcherrima* | 16.70 ± 1.65 | 8.35 ± 0.03 |
| *Combretum indicum* | 12 ± 1 | 4.19 ± 0.52 |

#### Experiment 1.3: Survival and insemination

##### Survival

Mosquito survival on day 5 varied among treatments (LRT *X^2^*_3_ = 48.18, P < 0.001, Fig. 4, Supp. Table 8), ranging from 82 ± 0.03% on 5% glucose, to 81 ± 0.03% on *I. coccinea*, 69 ± 0.03% on *C. pulcherrima,* and 54 ± 0.04% on *C. indicum*. All pairwise differences were significant except that between glucose 5% and *I. coccinea* (Supp. Table 9). There were no survival differences between species (*An. coluzzii*: 67 ± 0.03%, *An. gambiae*: 77 ± 0.02%, LRT *X^2^*_1_ = 1.75, P = 0.19, Fig. 4, Supp. Table 8), sex (male: 68 ± 0.02%, female: 75 ± 0.02%, LRT *X^2^*_1_ = 0.12, P = 0.72, Fig. 4, Supp. Table 8) and design (design 1: 64 ± 0.03%, design 2: 72 ± 0.03%, design 3: 80 ± 0.02%, LRT *X^2^*_1_ = 0.41, P = 0.52, Fig. 4, Supp. Table 8). The survival of males or females was improved when they were first maintained for 3 days on a 5% glucose solution (males design 2 and females design 3 in Fig. 4). In addition, such a maintenance on 5% glucose alleviated the differences caused by the treatments, resulting in a significant design by treatment interaction (LRT *X^2^*_3_ = 15.94, P = 0.001, Fig. 4, Supp. Table 8). All other interactions were non-significant (Supp. Table 8). The results from this experiment confirms the two previous survival assays: mosquito survival on *C. indicum* was the worst regardless of sex and species. Separate analyses and figures were also produced for each of the three designs (Supp. Table 8 and Supp. Fig. 8).

**Fig. 4:**
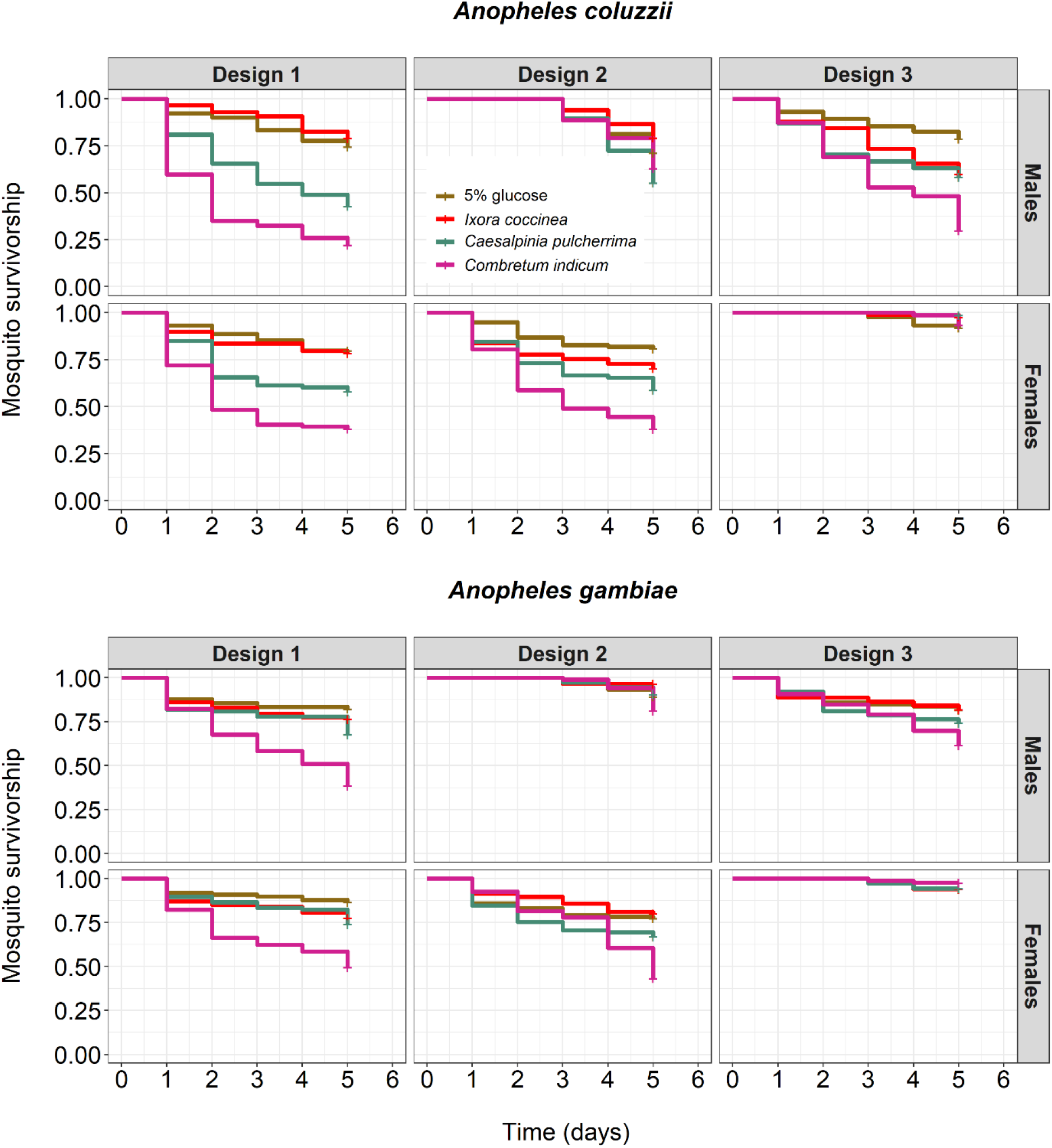
Effect of diet, species, sex, and design on mosquito survival over three replicates. Design 1: males and females were kept together for 5 days on one of four treatments, Design 2: males were first fed 5% glucose solution for 3 days before being introduced with the females maintained on the treatments. Design 3: females were first fed 5% glucose solution for 3 days before being introduced with males maintained on the treatments. Kaplan–Meier curves represent the proportion of live mosquitoes for each treatment from day 1 to day 5 post-emergence.

##### Insemination rate

Female insemination rate varied among treatments (LRT *X^2^*_3_ = 25.38, P < 0.001, Fig. 5, Supp. Table 10), with *C. indicum* causing the lowest insemination rate (71 ± 0.06%), followed by *C. pulcherrima* (73 ± 0.05%), *I. coccinea* (80 ± 0.04%) and 5% glucose (89 ± 0.03%) (Fig. 5, Supp. Table 11). All pairwise differences were significant except that between *C. pulcherrima* and *C. indicum* (Supp. Table 12). There were no differences in insemination rates between species (*An. coluzzii*: 79 ± 0.03%, *An. gambiae*: 80 ± 0.03%, LRT *X^2^*_1_ = 0.19, P = 0.67, Fig. 5, Supp. Tables 10, 11). There was no effect of design, *i.e.,* feeding the treatments to males exclusively for three days (design 3), females solely for three days (design 2), or both sexes simultaneously for three days (design 1) had no effect on mosquito insemination rates (design 1: 81 ± 0.04%, design 2: 82 ± 0.04%, design 3: 77 ± 0.04%, LRT *X^2^*_2_ = 3.97, P = 0.14, Fig. 5, Supp. Tables 10, 11). There were significant treatment by species (LRT *X^2^*_3_ = 15.58, P = 0.001, Fig. 5, Supp. Table 10), and treatment by design (LRT *X^2^*_6_ = 16.08, P = 0.01, Fig. 5, Supp. Table 10) interactions as well as a three-way interaction between treatment, species and design (LRT *X^2^*_6_ = 14.64, P = 0.02, Fig. 5, Supp. Table 10). Separate analyses and figures were also produced for each of the three designs separately (Supp. Table 10 and Supp. Fig. 9).

**Fig. 5:**
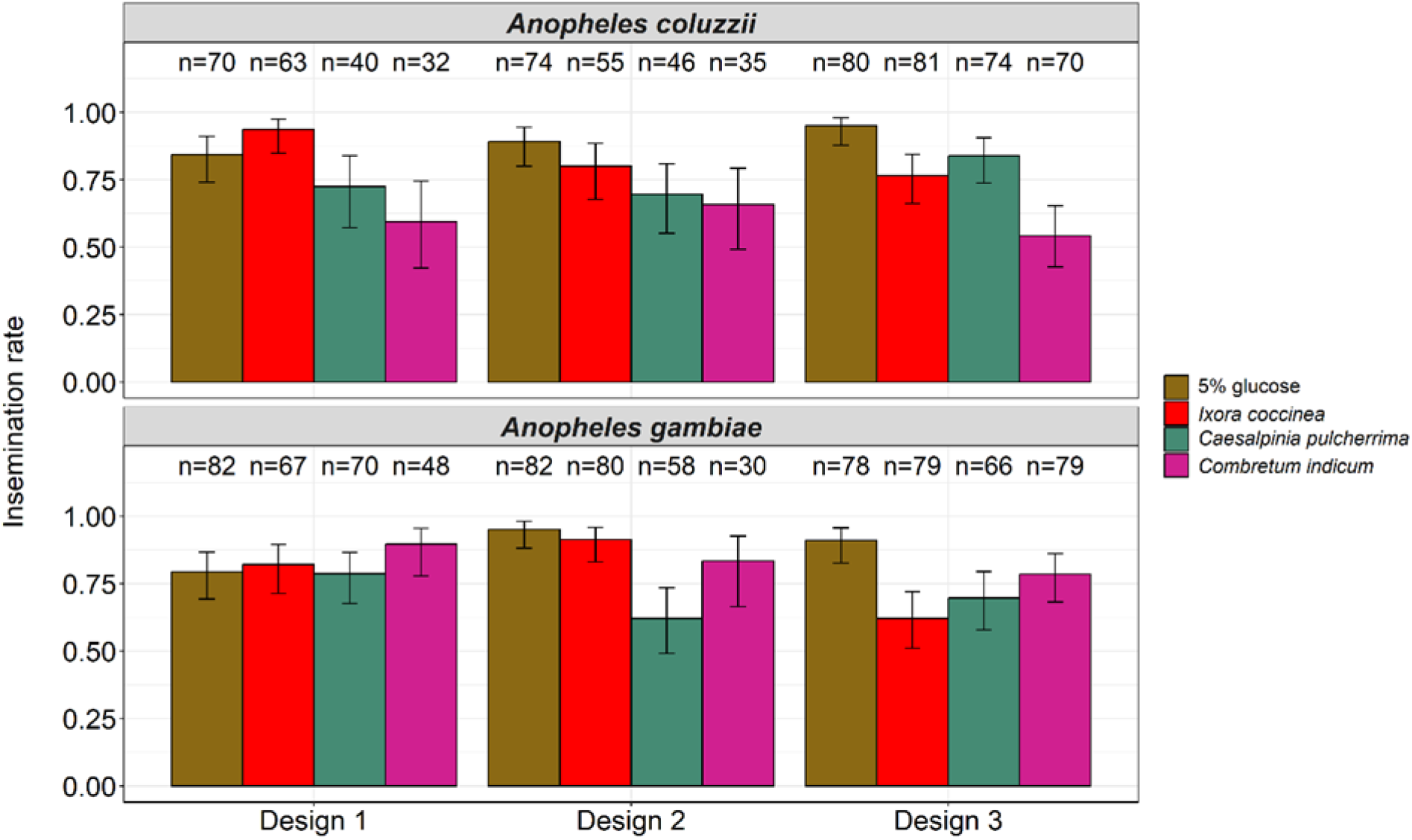
Effect of diet, species, and design on insemination rate over three replicates. Treatments were 5% glucose control solution, *I. coccinea, C. pulcherrima and C. indicum*. The numbers above the barplots represent the sample size for each treatment. The error bars represent the variability of data with 95% confidence interval.

### 2. Mosquito behavioral response to plant species

A total of 3,800 *An. coluzzii* and 4,576 *An. gambiae* were released on 64 occasions over 16 nights (4 release cages/night), and activation rate and plant relative attractiveness were measured.

#### Activation

Overall, the activation rate of *An. gambiae* (58% (0.57-0.59), with 2,653 of the 4,576 released *An. gambiae* flying into one of the five odor traps) was higher than that of *An. coluzzii* (43% (0.41-0.44), 1,625 of 3,800 released *An. coluzzii*) (LRT *X^2^*_1_ = 70.65, P < 0.001, Fig. 6a). Color marking increased mosquito activation (LRT *X^2^*_1_ = 80.68, P < 0.001, Supp. Fig. 10), regardless of mosquito species (*i.e.*, no species by color interaction, LRT *X*^2^ = 0.08, P = 0.78, Supp Fig. 10). There was no effect of density on mosquito activation (LRT *X^2^*_1_ = 2.68, P = 0.10). Supp. Fig. 11 shows mosquito activation rates for each of the 16 nights.

**Fig. 6:**
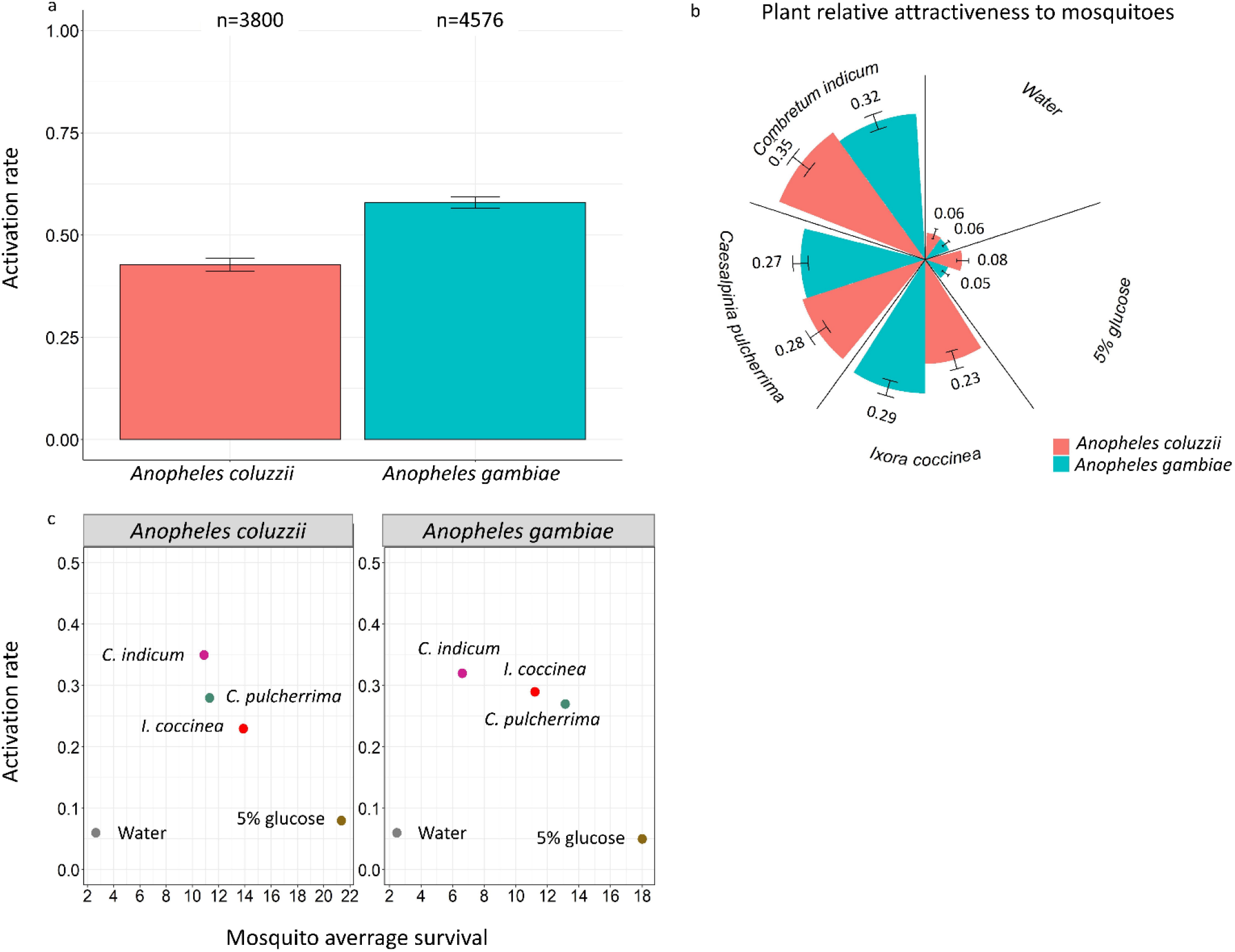
Mosquito behavioral response. (a) Activation rate of mosquitoes (*i.e.,* number of mosquitoes caught in all traps out of the total number of mosquitoes released). An average of 59.38 (±1.20) *An. coluzzii* and 71.5 (±0.91) *An. gambiae* females were released in one of four cages from 6 pm to 6 am. The numbers above the barplots represent the total number of released mosquito. (b) Plant relative attractiveness to mosquitoes (*i.e.,* number of mosquitoes caught in one trap out of the total number of mosquitoes caught in all traps). The numbers above the bars represent this attractiveness index. The error bars represent 95% confidence interval (±95% CI). The experiment was repeated over 16 nights leading to a total of 64 releases (4 cages x 16 nights). (c) Attractiveness index in relation to mosquito survival using mosquito average survival in experiment 1.2.1. The dots represent the treatments.

#### Attractiveness

Based on AIC, the minimum adequate model included species only as the main effect, *i.e.*, there was no influence of color marking, density, or interactions on plant relative attractiveness. In particular, *An. coluzzii* females exhibited a greater preference for the 5% glucose solution and a lesser preference for *I. coccinea* compared to their *An. gambiae* counterparts (Fig. 6b). Given the significant effect of mosquito species, separate analyses were conducted for each species to investigate their respective preferences among the five treatments. Table 3 shows the relative odds ratios between treatments for each mosquito species. First, the water and glucose treatments were equally attractive, but much less than the three plant species. Second, *C. indicum* was the most attractive plant to mosquitoes, with odds ratios of 6.6 and 6.8 for *An. coluzzii* and *An. gambiae*, respectively, compared to the control water treatment. Third, while the attractiveness of *C indicum* to mosquitoes was not significantly higher than that of *C. pulcherrima*, *An. coluzzii* showed a significant preference for *C. indicum* compared to *I. coccinea*. Supp. Fig. 12 shows the relative attractiveness of treatments for each of the 16 nights.

**Table 3:** Relative odds ratios of attractiveness between treatments for each mosquito species. **OR:** odds ratio. The relative odds ratios and 95% confidence interval were derived from the mixed-effects multinomial logistic regression model by exponentiating the regression coefficients.

|  | *Anopheles coluzzii* |  | *Anopheles gambiae* |  |
| --- | --- | --- | --- | --- |
|  | Relative OR (95% CI) | P-value | Relative OR (95% CI) | P-value |
| Water / 5% glucose | 1.17 (0.65-2.11) | 0.61 | 0.85 (0.52-1.38) | 0.50 |
| Water / *C. pulcherrima* | 5.22 (3.01-9.08) | < **0.001** | 5.17 (2.76-9.68) | < **0.001** |
| Water / *C. indicum* | 6.64 (3.86-11.41) | < **0.001** | 6.80 (3.67-12.60) | < **0.001** |
| Water / *I. coccinea* | 4.29 (2.55-7.23) | < **0.001** | 5.51 (3.21-9.46) | < **0.001** |
| 5% glucose / *C. pulcherrima* | 4 (2.46-6.48) | < **0.001** | 5.57 (3.00-10.35) | < **0.001** |
| 5% glucose / *C. indicum* | 5.09 (3.08-8.41) | < **0.001** | 7.27 (3.78-13.99) | < **0.001** |
| 5% glucose / *I. coccinea* | 3.27 (2.07-5.16) | < **0.001** | 5.91 (3.35-10.43) | < **0.001** |
| *C. pulcherrima* / *C. indicum* | 1.24 (0.83-1.85) | 0.30 | 1.27 (0.84-1.92) | 0.25 |
| *C. pulcherrima* / *I. coccinea* | 0.79 (0.53-1.18) | 0.25 | 1.04 (0.65-1.65) | 0.89 |
| *C. indicum* / *I. coccinea* | 0.64 (0.42-0.97) | **0.03** | 0.80 (0.51-1.24) | 0.31 |

Finally, we explored the association between mosquito preference and performance (using average mosquito survival as a proxy) by plotting the index of relative attractiveness against mosquito survival (Fig. 6c). Although the limited number of tested plant species precluded proper linear fitting, the relationship between plant attractiveness and mosquito survivorship tended to be negative, suggesting that *An. gambiae* and *An. coluzzii* did not prefer plant species that provided the best survivorship. This negative relationship between preference and performance was also obtained with our additional survival assay (Supp. Fig. 13 and 14).

## Discussion

Findings presented in this study do not support the hypothesis that females of the major malaria vectors *An. gambiae* and *An. coluzzii* exhibit significant preferences for the plant species that best supports their fitness, and that this could be mediated by both mosquito sugar intake and sugar content in plants. Rather, among the three species tested, *C. indicum*, the most attractive plant species to both mosquito species in the multiple-choice assays, had the lowest sugar content and provided the lowest mosquito survival, insemination rate and fructose positivity. Our results are inconsistent with the few previous studies suggesting that *An. gambiae* males and females are generally attracted to plants that provide abundant sugars, which in turn prolongs their survival ^31,37,38^. Instead, we observed a mismatch between mosquito performance and preference, and in this respect, our results align with the exception found in the studies of Manda et al. ^37^ and Nikbakhtzadeh et al. ^38^, wherein *P. hysterophorus* was found to be attractive despite providing little sugar and not extending mosquito survival. A first possible explanation for this mismatch is that *C. indicum* (and *P. hysterophorus*) are deceptive flowers. Studies have demonstrated that mosquitoes are drawn to floral semiochemicals, such as linalool oxides ^54,55^. The main components detected in the extracts of flowers of *C. indicum* are, in fact, linalool oxides ^56^, and it is plausible that *C. indicum* would release the right amount of such semiochemicals, making it highly attractive to mosquitoes. Furthermore, while the relative importance of visual vs chemical cues in our multiple-choice trap device is unknown, it is recognized that visual cues, including color and contrast, contribute to mosquito attraction, with lightly-colored flowers, particularly white ones, being typically appealing to mosquitoes ^57^. This may explain why *C. indicum* was an attractive species, given its unique characteristic of displaying a floral color change phenomenon ^58,59^. These flowers indeed create a striking contrast, both at the intra- and inter-flower levels, by transitioning gradually from white to pink and finally to red ^58,59^. In contrast, the flowers of *I. coccinea* and *C. pulcherrima* have darker and less contrasting colors. *I. coccinea* flowers have orange-red hue, while *C. pulcherrima* have red flowers with yellow margins ^60–62^. In the context of pollinator-plant interactions, the chemical and visual floral stimuli are typically used to signal the presence of a food source, communicating the suitability of a plant for insects, *i.e.,* act as so called ‘honest signals’ ^13^. However, in the case of deceptive plants, these stimuli are a false promise of a reward that the plants do not actually provide ^63,64^. Through intricate olfactory, visual, and tactile cues, pollinators are tricked by these rewardless flowers, which ultimately leave them empty-handed ^63,64^. A related scenario proposed by Nikbakhtzadeh et al. ^38^, in an effort to explain the paradox surrounding *P. hysterophorus*, would be that mosquitoes use the visual or olfactory cues to locate plant hosts, with no regard for the sugar content of the plants. This may be a reasonable assumption, as *C. indicum* (like *P. hysterophorus*) is an exotic species in Africa, allowing little time for *An. gambiae* to have evolved adaptive preferences for these introduced plants over local plants ^22,38^. In this study, the stimuli, including semiochemicals, involved in the attractiveness of *C. indicum* to mosquitoes was not determined. Further research is thus needed to characterize and isolate the mechanisms and stimuli contributing to floral preference. Besides fundamental interest, these mechanisms could be exploited to develop odor lures to trap or kill mosquitoes.

Secondly, it is possible that *C. indicum* is an ideal sugar source for some other insects, but due to specific flower morphologies (*e.g*., location of nectaries), accessing nectar may be especially challenging for mosquitoes, as previously demonstrated for hoverfly-plant interactions ^65^. This possibility, however, seems less plausible given that *C. indicum* displayed the lowest sugar content among the plants studied here, therefore suggesting that the low survival rates observed in mosquitoes fed on this species were likely due to the limited nectar production of the plant, rather than its nectar being inaccessible. However, the influence of sugar content alone may not fully account for the observed variations in survival among treatments. In particular, previous investigations on female specimens of *Aedes aegypti* and *Culex quinquefasciatus* suggest that amino acids present in nectar could enhance survival rates ^66,67^. In contrast, the presence of toxic secondary compounds in nectars^68^ might reduce mosquito survival ^25,69,70^. The chemical composition of the plant species utilized in this study has not been thoroughly characterized, and the presence or quantity of amino acids or secondary toxic compounds in nectars remains unknown. However, studies have revealed the presence of secondary compounds, such as terpenes, alkaloids, phenols, glycosides, and flavonoids in flower or leaf extracts of these plant species ^56,71–74^. It is uncertain whether, in our no-choice assays, the mosquitoes indeed consumed nectar rather than phloem or xylem fluids from tissue piercing, as previously observed ^32,75,22^. The possibility of mosquito tissue feeding and the existence of potentially toxic compounds in the plant fluids could partially explained why the 5% glucose solution, with an equivalent °Bx of 4.3, provided better mosquito survival compared to plants with flowers exhibiting higher °Bx (>10). Nonetheless, the most plausible explanation for the observed favorable survival on the 5% glucose solution remains, that despite its lower sugar concentration, it was readily accessible (via soaked cotton), highly fluid, and therefore easily ingested in unrestricted quantities.

Thirdly, it is important to note that a resource is not limited to food alone; it can also include resting sites, shelter, and mating partners, among other things. For instance, the enemy-free space hypothesis suggests that phytophagous insects may use certain host plants for refuge and defense against natural predators ^76,77^. Accordingly, our findings suggest that the plant preference of *An. gambiae* and *An. coluzzii* may be influenced by factors beyond nutritional quality, such as the availability of favorable resting sites, as is potentially the case with *C. indicum* in this study. In a previous investigation on the development of an attractive toxic sugar bait to target *An. arabiensis*, Tenywa et al ^78^ discovered that the most efficient bait prototype drew mosquitoes in mostly for resting purposes, rather than feeding only. Unfortunately, we did not perform anthrone tests on mosquitoes retrieved from odor traps during the multiple-choice assays to determine the proportion of fructose-positive mosquitoes. This would have helped distinguish whether plants were attractive for resting sites rather than for food sources, although the two might be linked.

Fourthly, we did not characterize the full range of mosquito fitness-related traits. The phenotypic traits measured here, *i.e.,* survival, insemination rate, and sugar intake, are not the only parameters determining mosquito fitness. For instance, it is possible, that while *C. indicum* provided low survival and insemination rate, it may provide other important nutrients, including vitamins, amino acids, and secondary compounds, which could improve other parameters of mosquito fitness, such as fecundity, or defense against pathogens. Research on various animals has revealed that different life-history traits can have distinct nutritional optima (^12^ and references therein). For example, in the flesh fly *Sarcophaga crassipalpis*, lifespan was optimized at a lower total carbohydrate concentration compared to that for egg production ^12^. Moreover, there is mounting evidence indicating that insects can use less nutritious food sources for purposes of self-medication ^79^. For instance, although the highly attractive *P. hysterophorus* ^32,34^ appears to be a poor food source ^37,38^, it contains parthenin ^70^, a toxin which has been shown to limit malaria parasite development within the mosquito ^80^. In this regard, it is particularly relevant to note that extracts from flowers of *C. indicum* have strong antimicrobial properties ^81,82^. Further studies should investigate the effect of this plant on a wider range of phenotypic traits, including mosquito egg production or defense against pathogens.

The design used for the performance experiments did not afford the mosquitoes the ability to select from multiple options as in a cafeteria design. We may expect different performance outcomes when mosquitoes would consume different proportions of each plant. In natural conditions, mosquitoes may visit *C. indicum* to acquire specific non-sugar nutrients, and subsequently forage on another plant species to obtain sugars. Furthermore, the nutritional needs of insects are dynamic, undergoing constant changes, influenced by factors such as age and physiological status, as well as temperature and other environmental variables ^83^. When considering these various pieces of information collectively, it becomes apparent that the plant preference pattern of mosquitoes, as observed in our laboratory setting, may not only diverge but also serve as an indication that the preference exhibited for *C. indicum* is not necessarily maladaptive in natural environments.

The effects of the various treatments (*I. coccinea, C. pulcherrima, C. indicum*, and the 5% glucose solution) on mosquito survival were not consistent across our three tests (experiments 1.1, 1.2, and 1.3). For instance, in the first and third tests, *I. coccinea* provided similar survival rates to that of the 5% glucose solution, while in the second test, survival on *I. coccinea* was lower compared to that achieved with the 5% glucose solution. Similarly, while in experiments 1.1 and 1.3 (as well as in an additional survival experiment presented in Supp. Fig. 13), the survival of mosquitoes fed with *C. indicum* was similar to that of mosquitoes fed with water only, *C. indicum* provided much better survival than water in experiment 1.2. It is important to note that these experiments were conducted at different time periods using distinct groups of mosquitoes and plant individuals/populations, leading to variability in both insect and plant materials. Furthermore, we utilized plants collected from the field, which were naturally exposed to a whole community of nectar-feeding animals. Therefore, it is plausible that the nectar abundance could have fluctuated depending on the collection period (phenology of the plant) or the identity of the flowers collected, leading to variation in plant-mediated effects on mosquito survival. This perspective offers another possibility for elucidating the mismatch between performance and preference observed in our experiments: in natural settings, *C. indicum* flowers might be highly attractive (as observed in our multiple-choice laboratory assays), and entice a multitude of consumers (including wild mosquitoes), depleting its nectar and leaving our mosquitoes with nothing during the no-choice experiments. A suggestion for future studies is to use mesh bags to protect the flowers from nectar depletion by other nectar-feeding animals.

Our behavioral assay unveiled a heightened responsiveness of *An. gambiae* towards plants in comparison to *An. coluzzii*, leading to a greater activation rate. The underlying reasons remain ambiguous, but intrinsic genetic factors may play a role. Prior research conducted on male swarms of *An. coluzzii* from the Vallée du Kou and *An. gambiae* from Soumousso (the two locations from which our mosquito colonies were derived), demonstrated that swarming An. gambiae males possessed a higher total sugar reserve compared to their *An. coluzzii* counterparts ^87^. This observation suggests a potentially innate predisposition for greater sugar-feeding tendencies in *An. gambiae* relative to *An. coluzzii*, although ecological differences between the two sites may also account for this discrepancy and our cold-anthrone test suggested equivalent levels of sugar positivity between the two species.

In previous studies, *Anopheles* preference for plants has been assessed using either a dual-port Y olfactometer ^31,34^ or by direct observations of mosquitoes perching or feeding on plants ^32^. The novel multiple-choice test device employed in this study, inspired by the odor-baited net traps utilized to measure mosquito preference for vertebrate hosts ^88,89^, demonstrated remarkable reliability in assessing the behavioral response of mosquitoes towards different plant species. Notably, the two control traps baited with either water or 5% glucose solution, which do not emit semiochemicals, yielded very few mosquito captures. This successful validation of the device eliminates the need for human observation or video recording techniques and enables comparisons across a larger number of plants.

## Conclusions

*Anopheles gambiae* and *An. coluzzii* encounter diverse plant communities in their environment, providing them an opportunity to feed selectively on a range of plant species that can play crucial roles in their life history. Consistent with prior research, our findings revealed that different plant species elicited varying levels of mosquito survival in no-choice assays. However, contrary to our initial hypothesis, which posited that mosquitoes would exhibit a preference for certain species based on perceived differences in resource quality, we observed highest preference for the plant species that resulted in the lowest survival rates. This intriguing finding suggests the existence of alternative mechanisms influencing *Anopheles* preference for plants. The exploration of mosquito-plant relationships has long been overlooked, but there is immense potential for future research to build upon the foundational knowledge established in other insect systems, such as *Drosophila* flies, *Spodoptera* moth caterpillars, or locusts. The exploration of the aforementioned questions (*e.g*., do mosquitoes solely seek food (sugar) when selecting plants?; How do plants influence a range of specific mosquito fitness traits in varying ways?; and how does plant quality vary across species/individuals and changing environments?) can greatly benefit from adopting a nutritional ecology framework ^90^. This approach may provide a comprehensive understanding of the meaning of plant quality for mosquito performance and the extent to which mosquitoes are able to actively regulate their nutrition through plasticity in behavioral preference. Enhancing our comprehension of mosquito-plant interactions can also be pivotal for the improvement of attractive toxic sugar bait strategies and for the advancement of synthetic plant odor lures used in malaria vector control efforts in Africa.

## Acknowledgements

This work was supported by the ANR Grant STORM No.16-CE35-0007 and the IRD JEAI Grant No. AAP2018_JEAI_PALUNEC. We are grateful to NIGNAN Saibou from the Institut de Recherche pour le Développement and Dr. OUOBA Y. Hermann from the University Joseph KI-ZERBO/University Center of Ziniaré, Burkina Faso for plant species identification. We would like to thank the biochemistry and microbiology laboratory of the Département Technologie Alimentaire (DTA) of the Institut de Recherche en Sciences Appliquées et Technologies (IRSAT) of Bobo-Dioulasso for technical support, particularly Dr. Abel Tankoano, Hamadou Rouamba and Inessa Ouattara for their assistance in determining the degree Brix and total sugar content of the flowers. Thanks to all of the IRSS/DRO technicians and students, notably to Yasmire R. M. Gnanou, Felicité W. Nikiéma and Christian P. Younga, for their help during the experiments.

## Author contributions

P.S.L.P., D.F.D.S.H., O.G. and T.L. conceived and designed the study. P.S.L.P., Y.M. and D.F.D.S.H. conducted the experiments. P.S.L.P. and T.L. analysed the data. P.S.L.P., D.F.D.S.H., O.G. and T.L. drafted the manuscript. D.F.D.S.H., O.G. and T.L. supervised the study. R.S.Y., A.C., D.C., L-C.G., A.D. and R.K.D. read, revised and approved the final manuscript.

## Additional information

**Supplementary Information**

