## Supplemental Data 1 for "The paradox of plant preference: the malaria vectors *Anopheles gambiae* and *Anopheles coluzzii* select suboptimal food sources for their survival and reproduction"

**Supplementary figures**

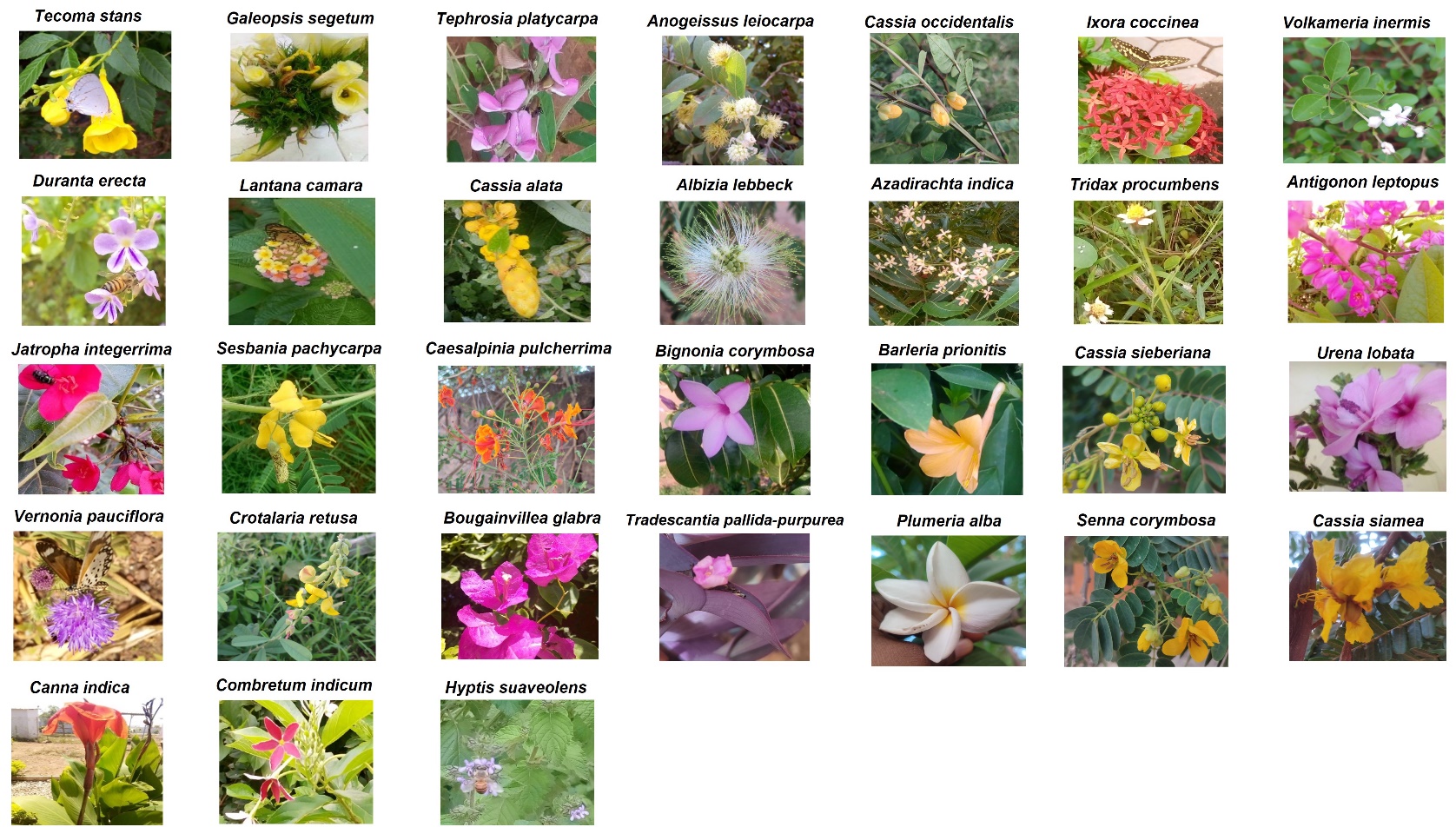

**Supplementary Fig. 1**: Photos of the 31 species of plant flowers screened, with their scientific names. These plant flowers were collected in the city of Bobo Dioulasso the village of Farako-ba. The species are ranked according to their effect on survival from positive to negative (see figure 1).

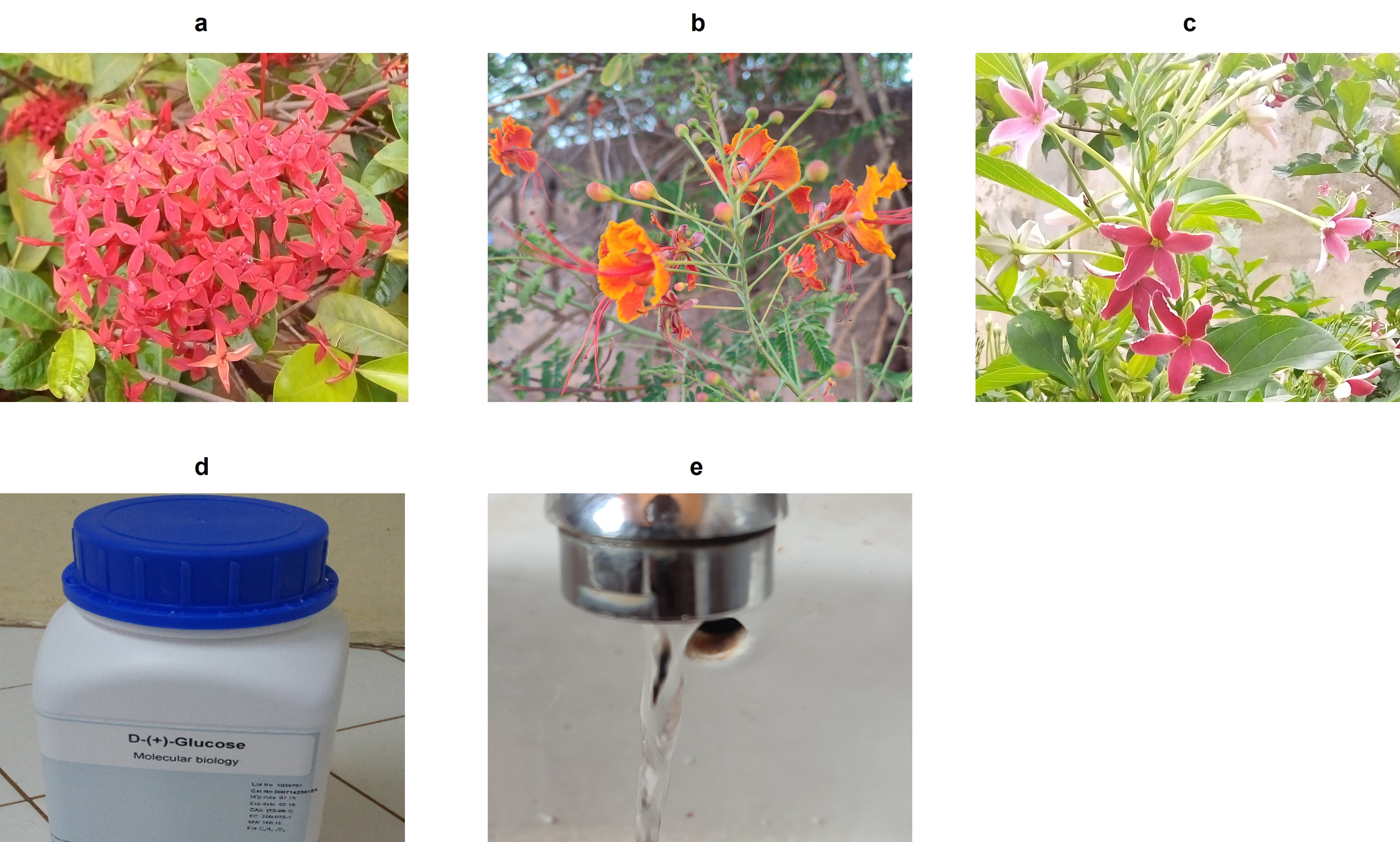

**Supplementary Fig. 2**: The five diet treatments used as part of experiments 1.2, 1.3 and 2. (a) *I. coccinea*, (b) *Caesalpinia pulcherrima*, (c) *Combretum indicum*, (d) Glucose (positive control), (e) Water (negative control).

**Anthrone test**

This test detects the presence of fructose in mosquitoes ^1^. The anthrone solution was prepared by dissolving 150 mg of anthrone powder in 100 ml of 68.11% sulphuric acid and stored at +4°C in the refrigerator. Mosquitoes were individually crushed in Eppendorf tubes containing 0.5 ml of the prepared anthrone solution. The whole was incubated for 60 min at room temperature for reaction. The lemon-yellow anthrone solution reacted with fructose to give light green, blue or dark blue colors depending on the amount of fructose ingested by the mosquitoes (Supp. Fig. 3).

**Supplementary Fig. 3**: The anthrone test of experiment 1.2. (a) Mosquitoes in a 1.5 ml Eppendorf tube ready for grinding. (b) Reaction obtained after 1 h of incubation.
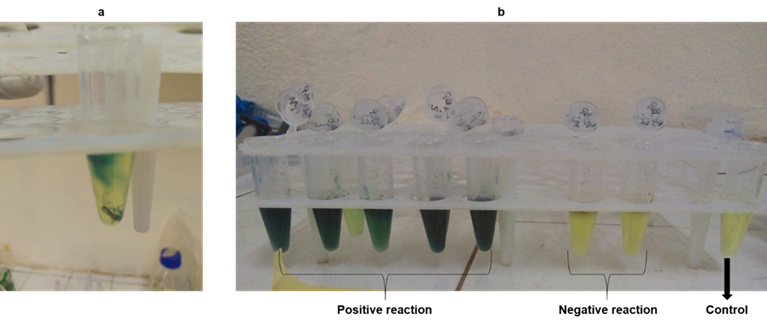

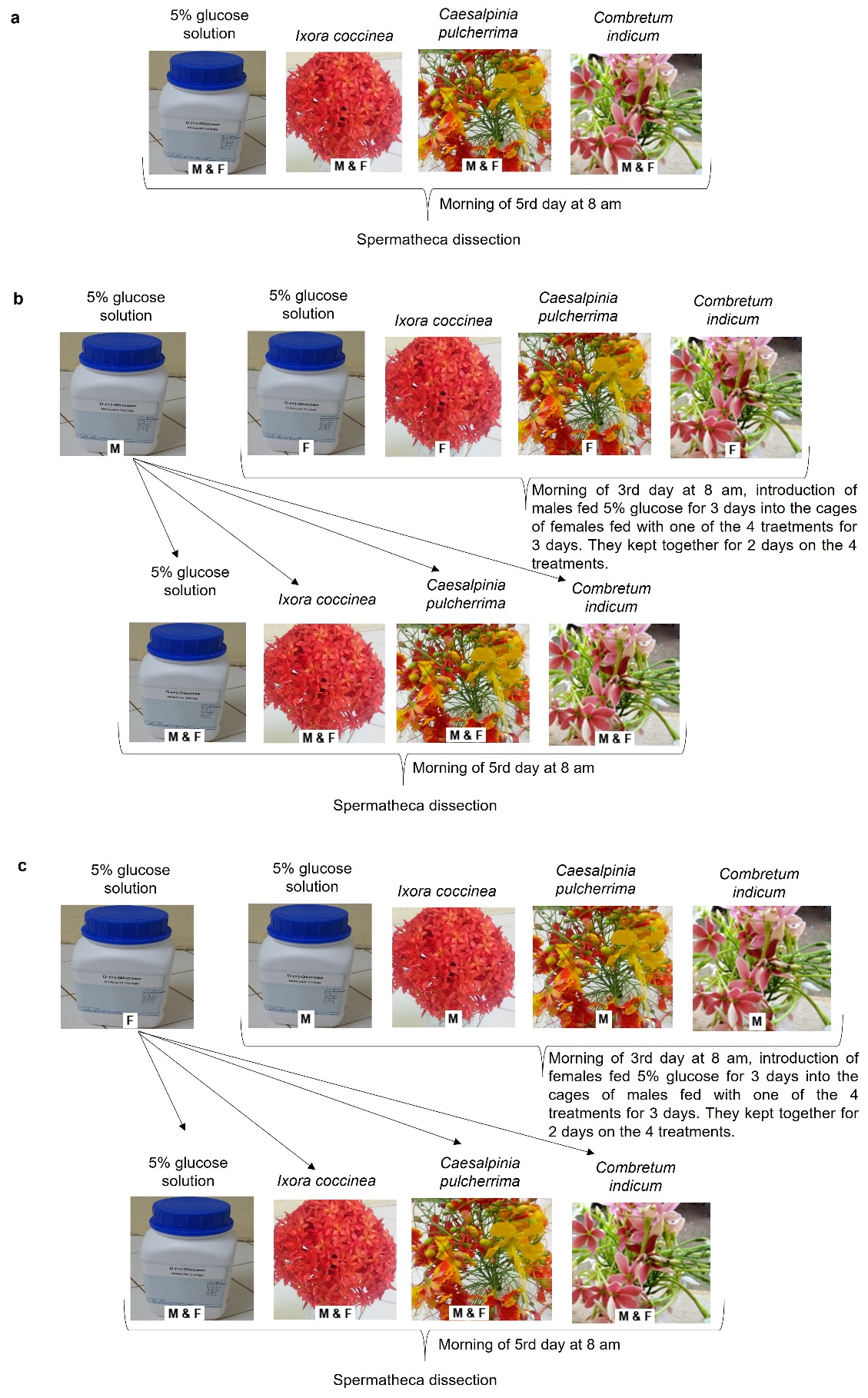

**Supplementary Fig. 4**: Schematic description of the three designs used for the insemination and survival experiment 1.3. (a) Design 1: males and females were kept together for 5 days on one of four diet treatments (5% glucose control solution, *I. coccinea, C. pulcherrima* and *C. indicum*), (b) Design 2: males were fed with 5% glucose solution for 3 days before being introduced into the cages of the females which were kept on the diet treatment for 3 days. The two sexes were kept together for 2 days on plant treatments. Design 3: females were fed with 5% glucose solution for 3 days before being introduced into the cages of the males which were kept on the diet treatment for 3 days. The two sexes were kept together for 2 days on diet treatments.

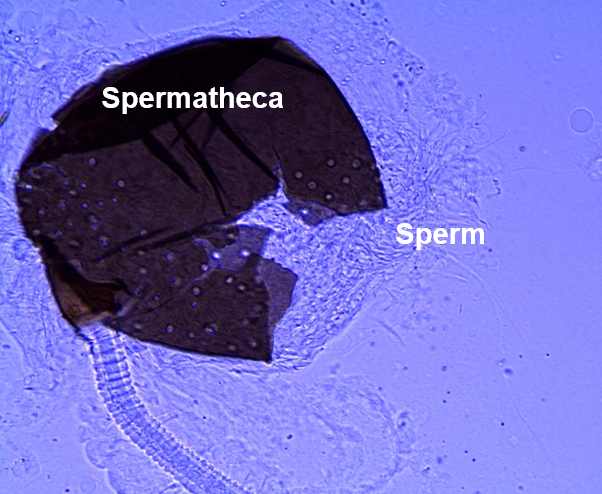

**Supplementary Fig. 5**: Inseminated spermatheca releasing its contents

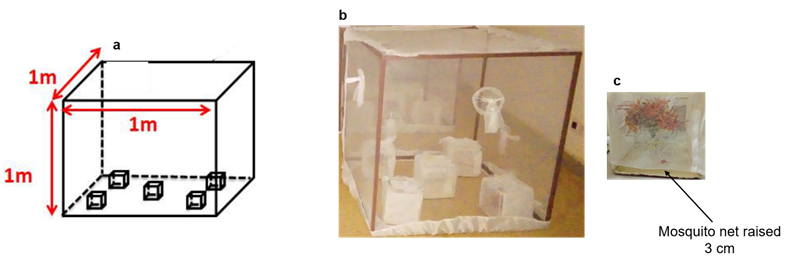

**Supplementary Fig. 6**: Behavioral device used to study mosquito preference for plant treatment. (a) Schematic representation of the device. (b) Photo of one of the large releasing cages housing five smaller treatment-baited traps. (c) Photo of an odor trap with the mosquito net raised to 3 cm above the ground.

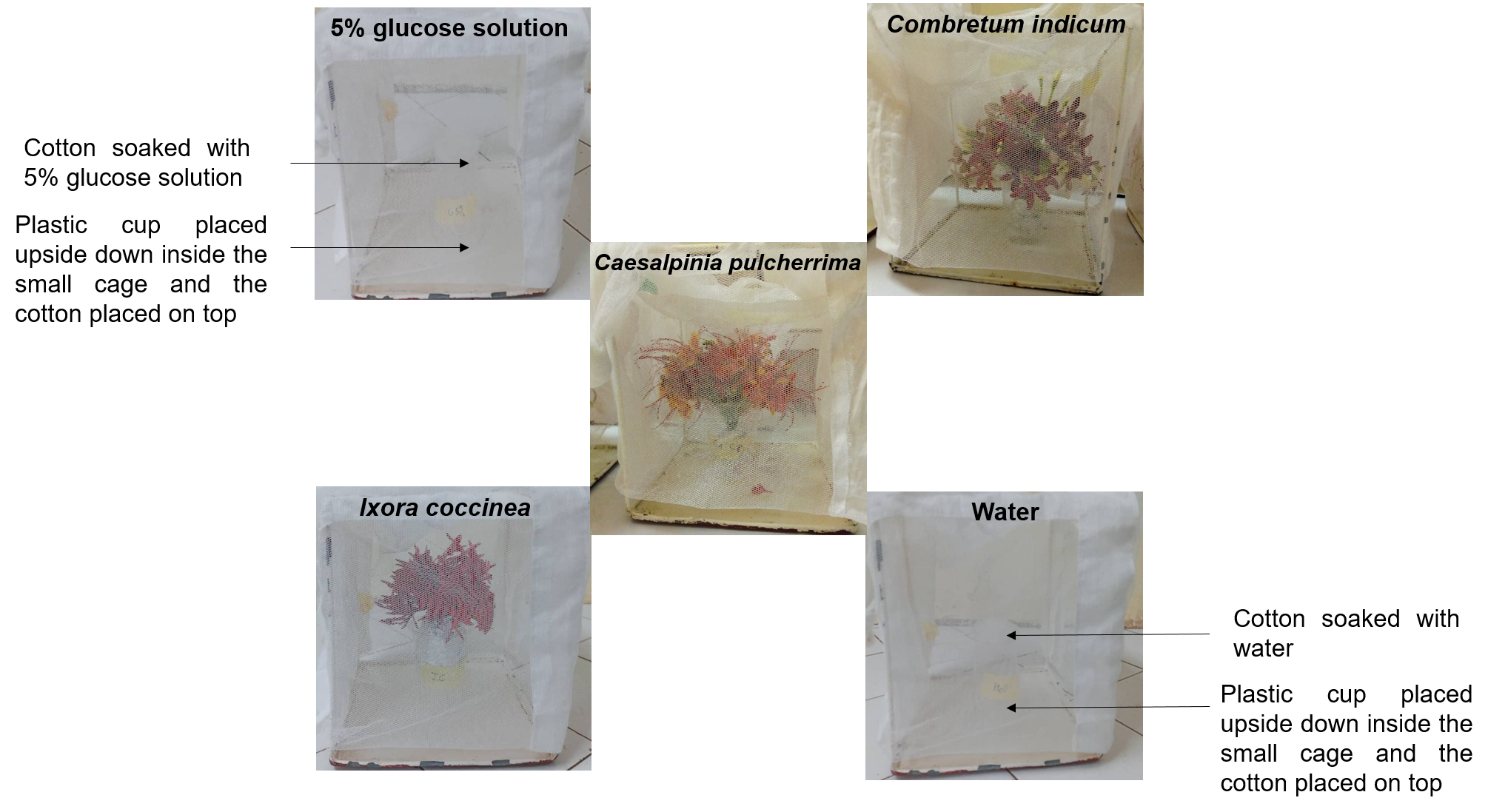

**Supplementary Fig. 7**: Traps baited with one of the five treatments: 5% glucose solution, water, *I. coccinea, C. pulcherrima* and *C. indicum*. The position of the traps was randomized among the cages and the replicates (night tests).

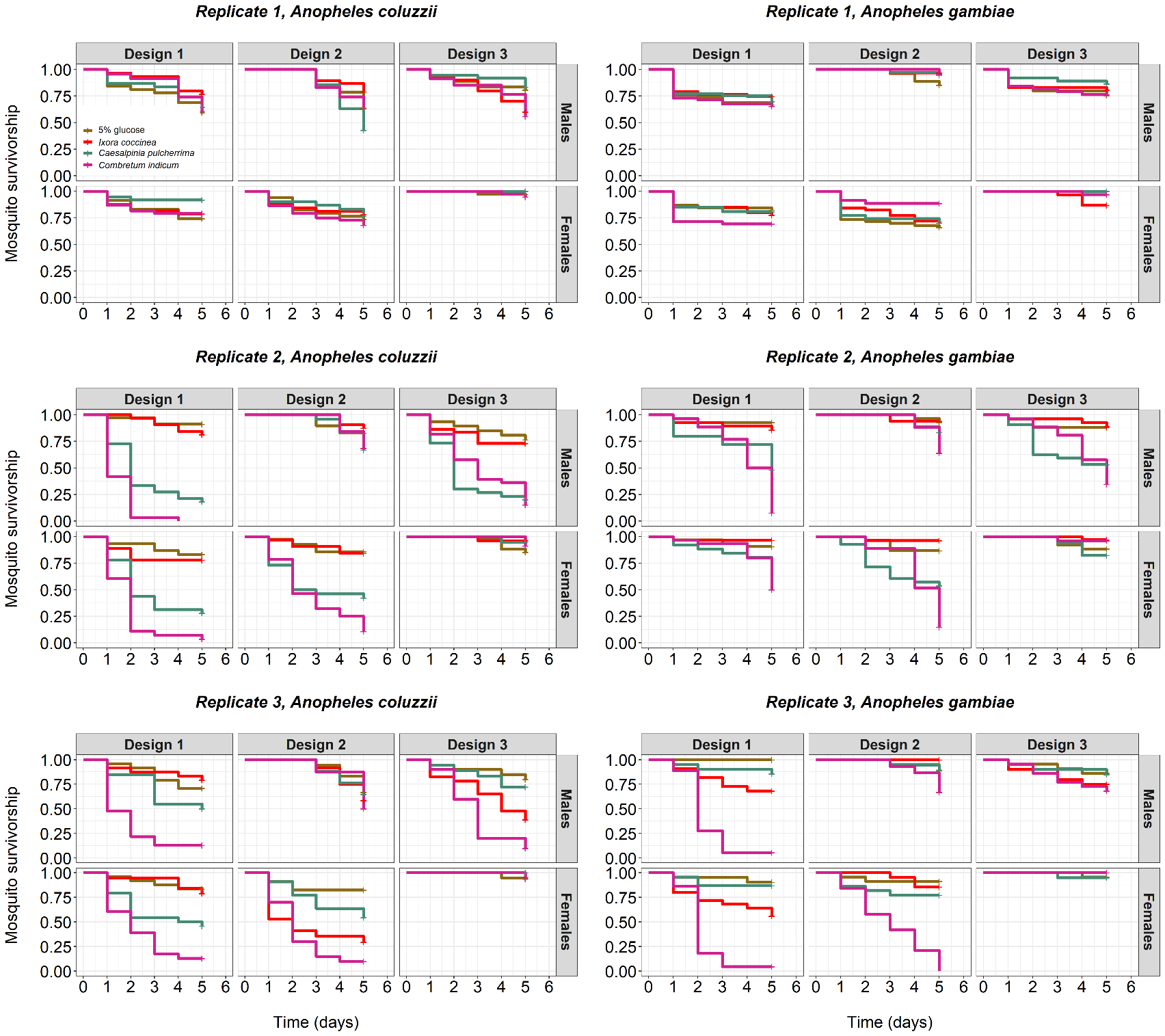
 **Supplementary Fig. 8**: Effect of diet, species, sex, and design on mosquito survival for each replicate. Design 1: males and females kept together for 5 days on different diet treatments, Design 2: males were fed with 5% glucose solution for 3 days before being introduced into the cages of the females, which were kept on the treatments for 3 days. The two sexes were kept together for 2 days on the diet treatments. Design 3: females were fed with 5% glucose solution for 3 days before being introduced into the cages of the males, which were kept on the diet treatment for 3 days. The two sexes were kept together for 2 days on diet treatments. Diet treatments were 5% glucose control solution, *I. coccinea, C. pulcherrima and C. indicum.* Kaplan–Meier curves represent the proportion of live mosquitoes over time for each diet treatment. Mosquito survival was recorded from day 1 to day 5 post-emergence

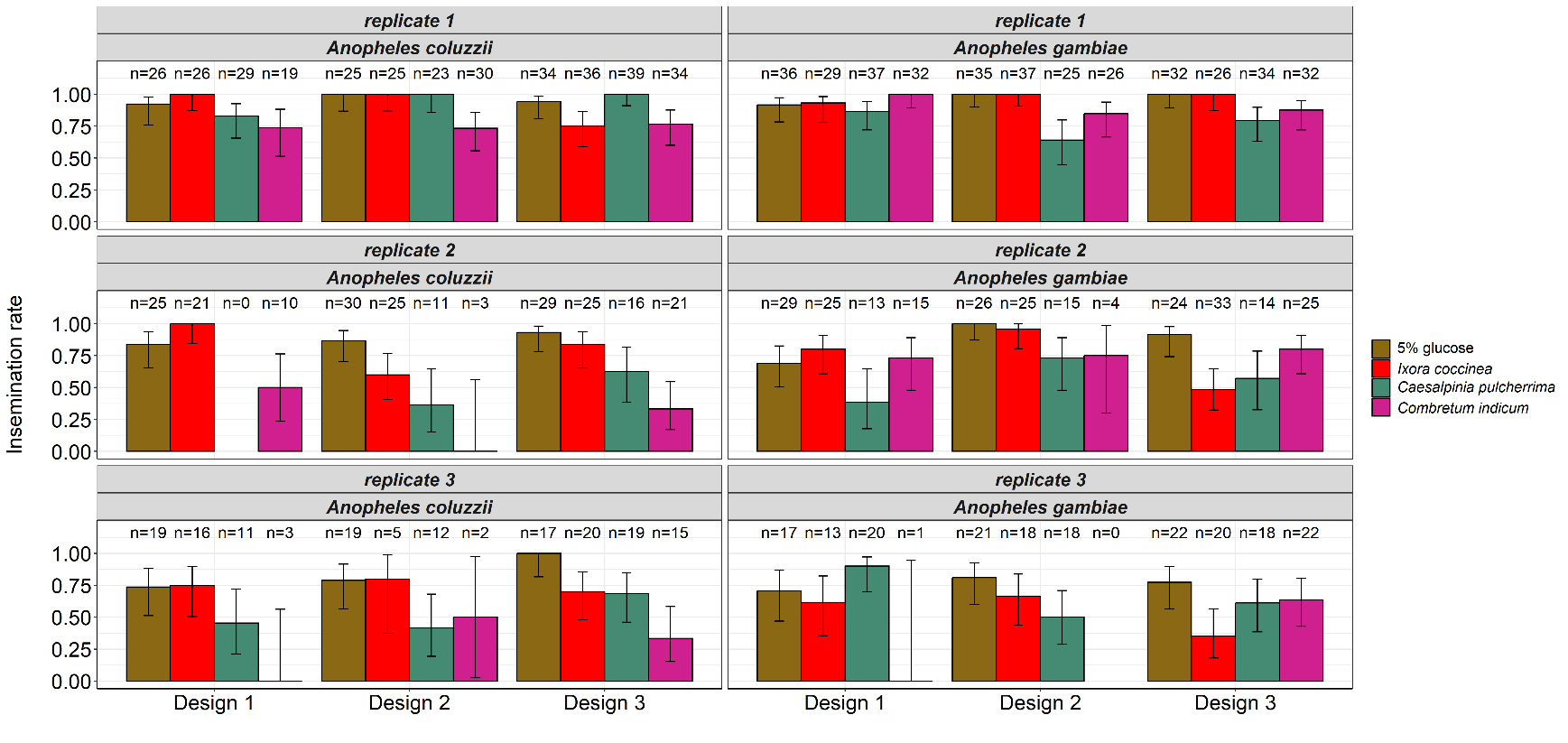
**Supplementary Fig. 9**: Effect of diet, species, and design on insemination rate for each replicate. Design 1: males and females kept together for 5 days on different diet treatment, Design: males were fed with 5% glucose solution for 3 days before being introduced into the cages of the females, which were kept on the diet treatment for 2 days, and Design 3: females were fed with 5% glucose solution for 3 days before being introduced into the cages of the males, which were kept on the diet treatment for 2 days. Diet treatments were 5% glucose control solution, *I. coccinea, C. pulcherrima* and *C. indicum*. The numbers above the barplots represent the sample size for each diet treatment. The error bars represent the variability of data with 95% confidence interval.

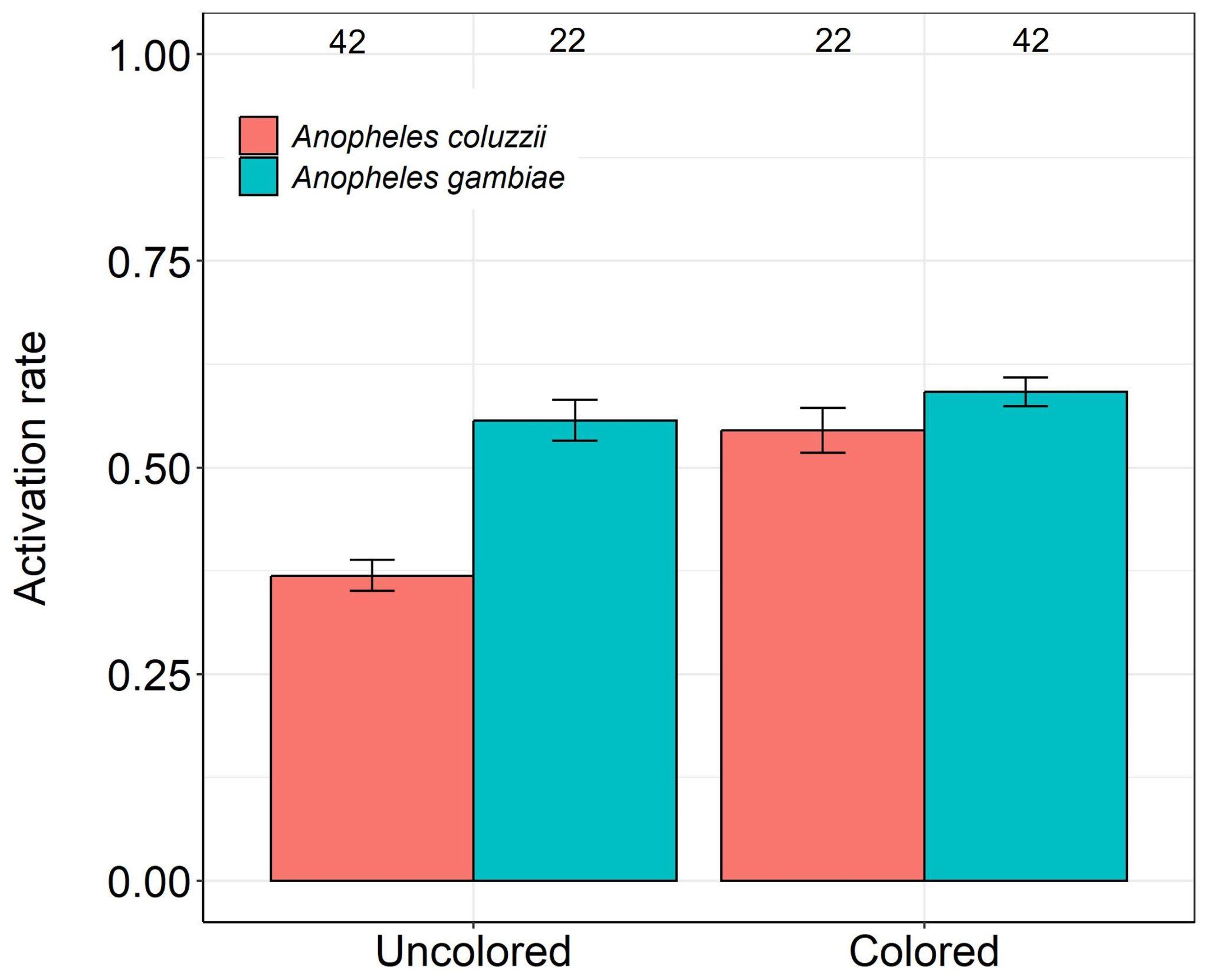

**Supplementary Fig. 10**: Effect of color on mosquito activation rate for each species. The numbers above the barplots represent the number of mosquito releases. The error bars represent the variability of data with 95% confidence interval.

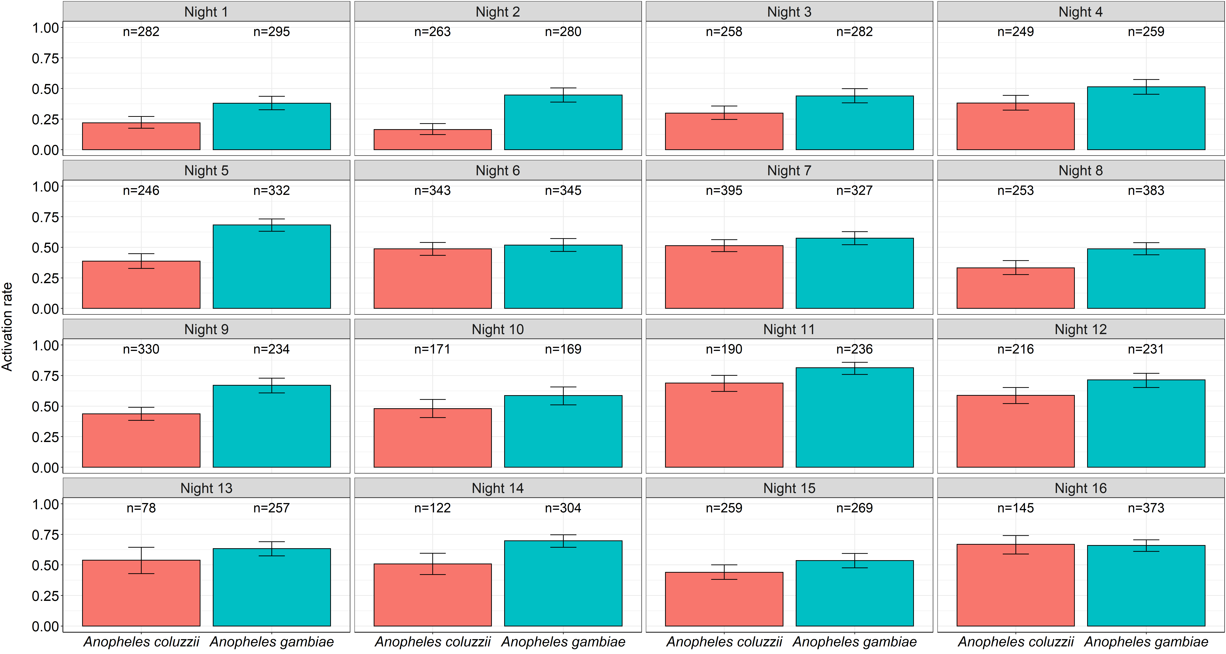
**Supplementary Fig. 11**: Effect of species on mosquito activation rate (number of mosquitoes caught in all traps out of the total number of mosquitoes released) for each night. The numbers above the barplots represent the sample size for each species. The error bars represent the variability of data with 95% confidence interval.

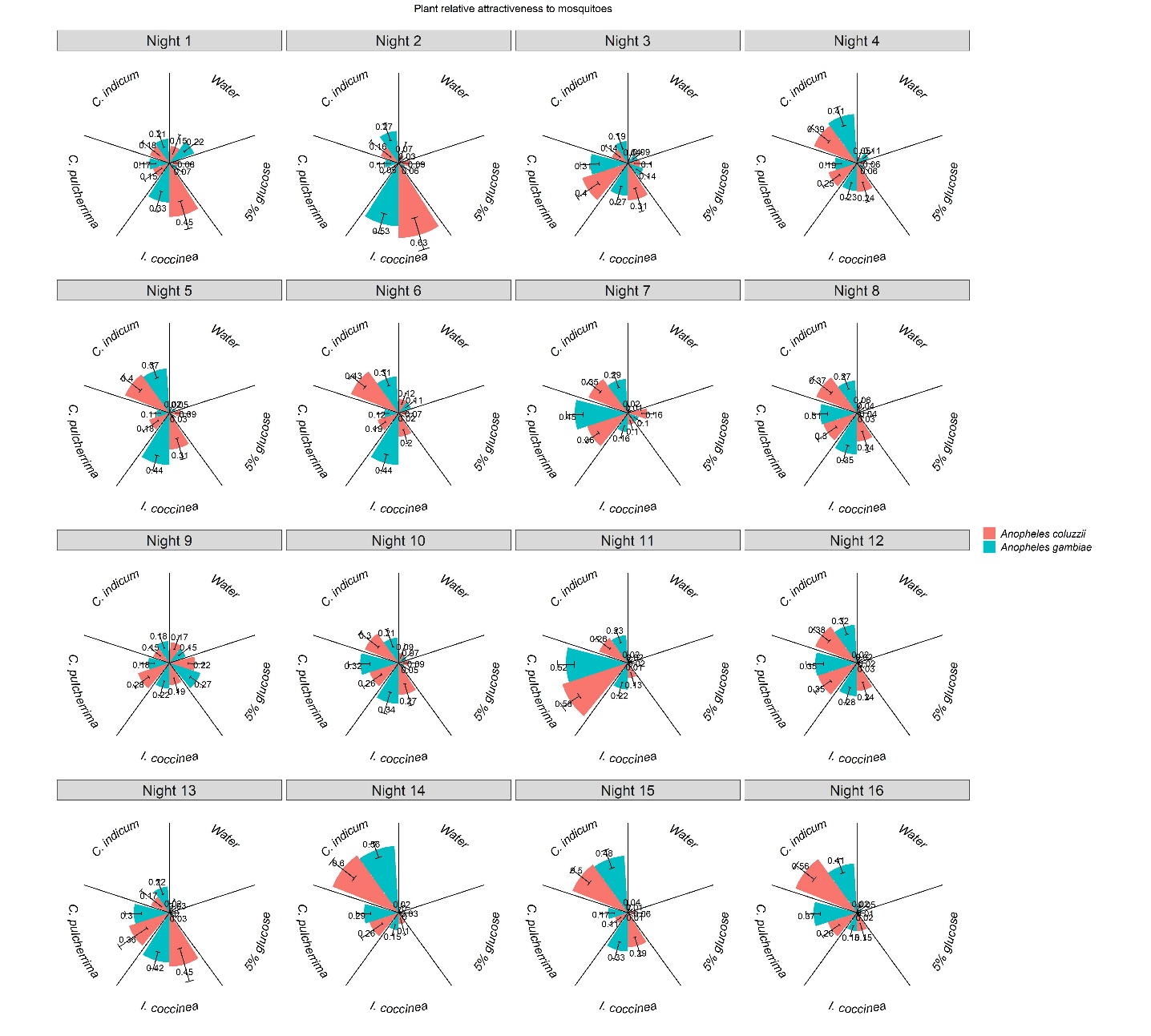

**Supplementary Fig. 12**: Odor treatment (five levels: water, 5% glucose, *I. coccinea, C. pulcherrima* and *C. indicum*) relative attractiveness to *Anopheles coluzzii* and *Anopheles gambiae* for each night. Relative attractiveness is the number of mosquitoes caught per trap out of the total number of mosquitoes caught in all traps. The numbers above the barplots represent the proportion of mosquitoes captured in each trap and for each species of mosquito. The error bars represent the variability of data with 95% confidence interval.

**Additional survival assay**

**Method**: Both *An. coluzzii* and *An. gambiae* were used, and males and females were distinguished in this experiment. Upon emergence, males and females of *An. gambiae* (sample sizes are given in the legend of the corresponding figure below) were placed together in 20 cm × 20 cm × 20 cm cages and maintained on one of five treatments: *I. coccinea, C. pulcherrima, C. indicum*, water (negative control) and 5% glucose solution (positive control) (Supp. Fig.2). Mosquitoes were exposed to these treatments in the same manner and timing as in the screening experiment 1.1, and survival test (Experiment 1.2). The mortality of males and females was monitored every day between 4 pm and 5 pm until all mosquitoes were dead.

Cox’s model with censoring was used to test the effect of treatment (5 levels: water, 5% glucose, *I. coccinea, C. pulcherrima, C. indicum*), mosquito species (2 levels: *An. coluzzii* and *An. gambiae*), sex (2 levels) and their interactions on mosquito survivorship.

**Results**: There were no main effects of species (*An. coluzzii*: 31 ± 0.09%, *An. gambiae*: 41 ± 0.08%, LRT *X^2^*_1_ = 0.36; P = 0.55, Supp. Fig. 13, Supp. Tables 13, 14) and sex (male: 38 ± 0.09%, female: 34 ± 0.08%, LRT *X^2^*_1_ = 0.58; P = 0.45, Supp. Fig. 13, Supp. Tables 13, 14) on mosquito survival. However, a significant three-way interaction was observed between treatment, species, and sex, indicating that the effects of treatment on survival varied depending on the mosquito species and sex. Specifically, *C. pulcherrima* was found to provide better survival than *I. coccinea* for *An. gambiae* males, but the opposite pattern was observed for *An. coluzzii* males. In contrast, both plant species provided similar survival rates for females of both mosquito species. In the absence of a food source, *i.e.,* water only, mosquitoes died within three days, with the exception of *An. coluzzii* males, which were dead within four days (Supp. Fig. 4). Survival rates varied among treatments (LRT *X^2^*_4_ = 170.39; P < 0.001, Supp. Fig. 13, Supp. Table 13), and all pairwise differences were significant except that between *I. coccinea* and *C. pulcherrima* (Supp. Table 15). In particular, mosquitoes were 75 times more likely to survive when kept on a 5% glucose solution rather than on water alone (Supp. Table 16). Compared to the water control, the chances of survival were 9 and 10 times greater when mosquitoes were maintained on *C. pulcherrima* and *I. coccinea*, respectively (Supp. Table 16). In contrast, *C. indicum* slightly improved mosquito survival (1.46 times) compared to the water control (Supp. Table 16).

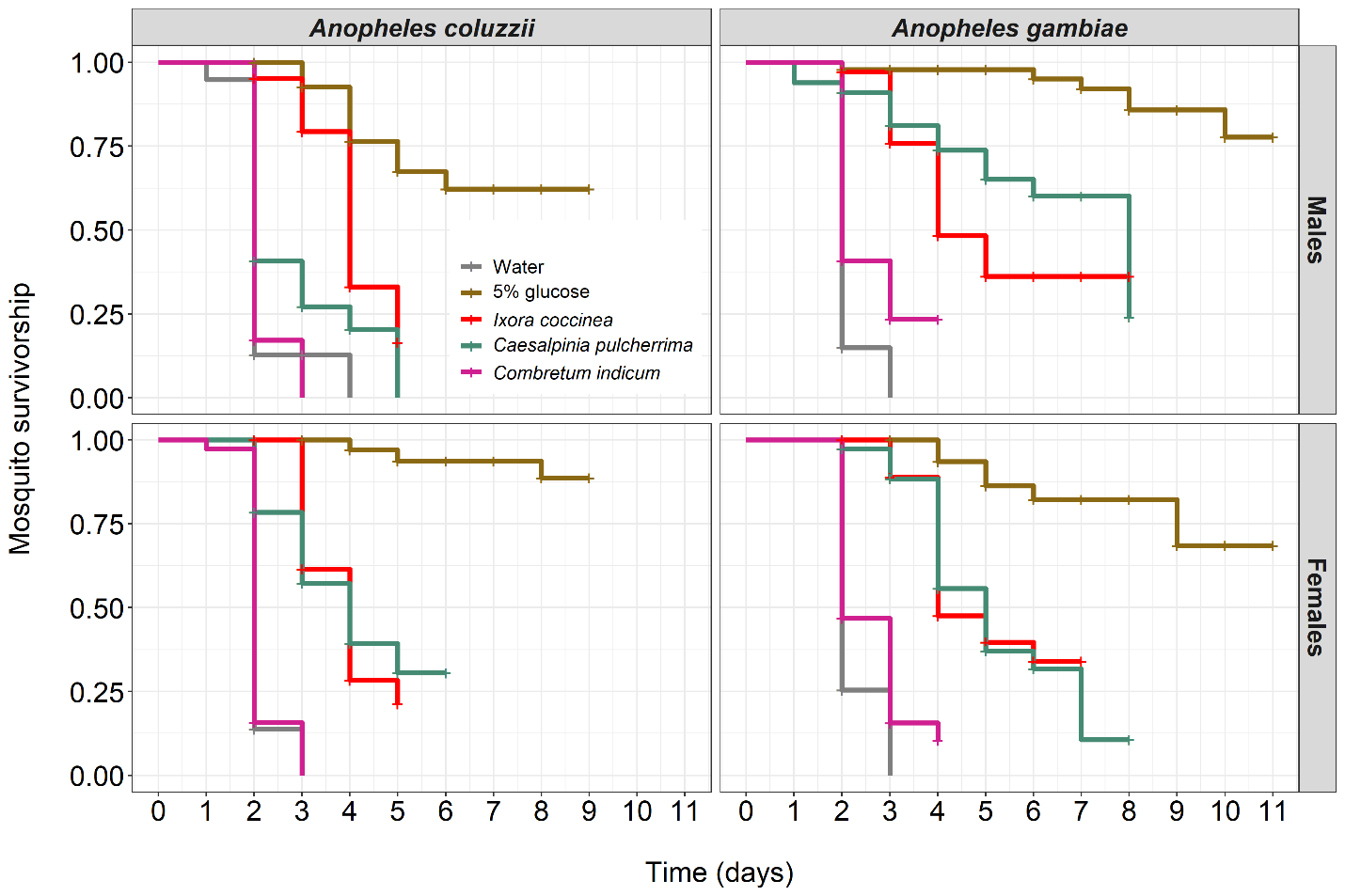

**Supplementary Fig. 13:** Effect of diet on the survival of *Anopheles coluzzii* and *Anopheles gambiae* according to sex. Kaplan–Meier curves represent the proportion of live mosquitoes over time for each diet (water = negative control, 5% glucose solution = positive control). Between 21-39 emerging males (mean ± se: 30.2 ± 3.14, median: 29) and 29-51 emerging females (mean ± se: 38.6 ± 3.53, median: 38) of *An. coluzzii*, and between 22-46 emerging males (mean ± se: 35.2 ± 3.99, median: 35) and 32-59 females (mean ± se: 40.8 ± 4.69, median: 37) of *An. gambiae* were placed together and maintained on one of the five diets. Some live male and female individuals of both species were randomly collected daily from the cages for the cold anthrone test to confirm fructose intake. The survival of males and females *An. coluzzii* and *An. gambiae* were monitored until all mosquitoes were dead.

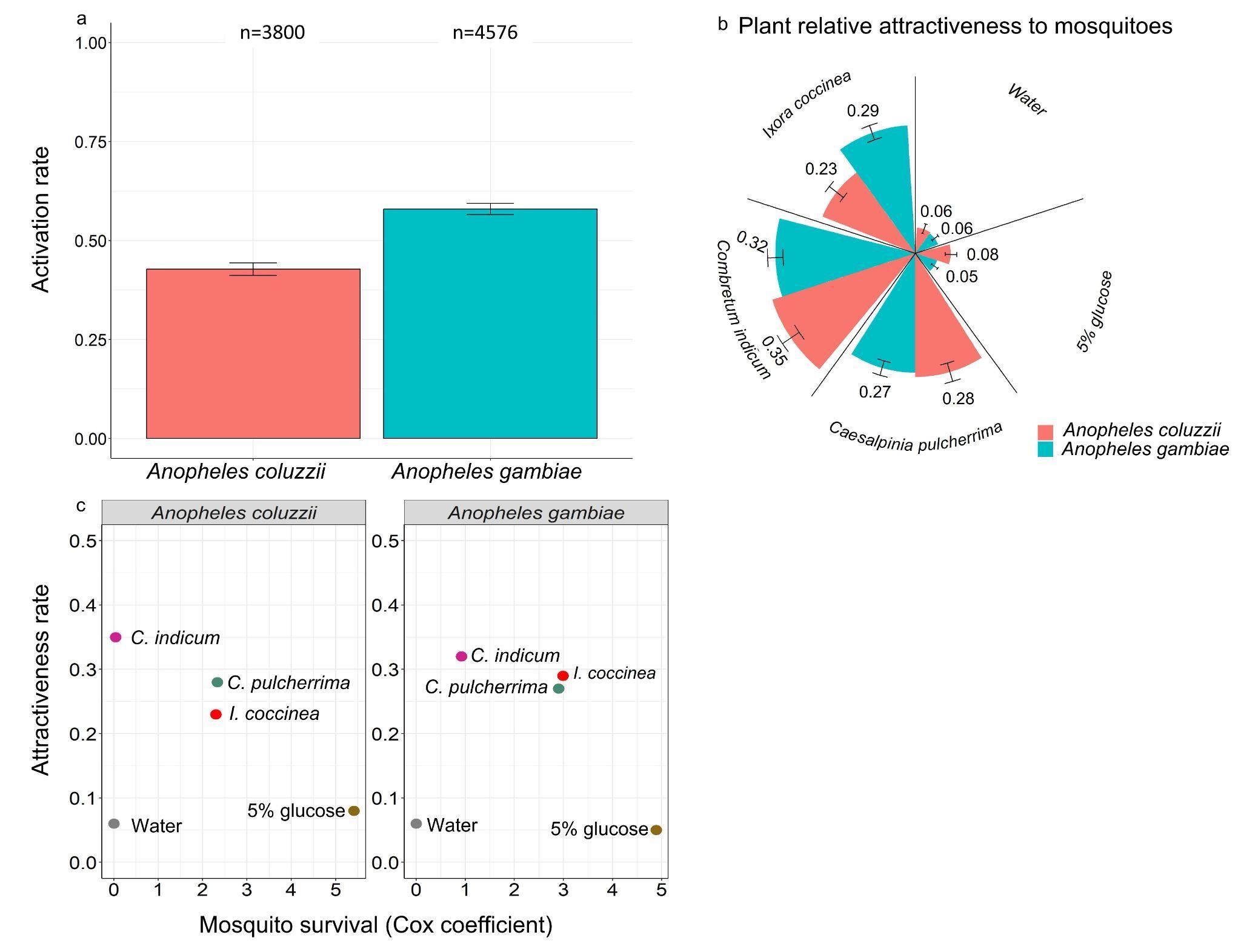

**Supplementary Figure 14**: Attractiveness index in relation to mosquito survival using coefficient of the Cox model in above experiment. The dots represent the plant treatment.
