## Supplemental Data 2 for "The paradox of plant preference: the malaria vectors *Anopheles gambiae* and *Anopheles coluzzii* select suboptimal food sources for their survival and reproduction"

**Supplementary tables**

**Supplementary Table 1:** Survival rate of *Anopheles coluzzii* (±95CI) maintained on one of 31 plant species and a 5% glucose solution (Experiment 1.1). The species are ranked according to their effect on survival from positive to negative (see figure 1). The values represent survival rates on day 6 post-emergence, when mortality monitoring was stopped.

| Diet | Family-subfamily of plants | Survival rate (±95% CI) |
| --- | --- | --- |
| 5% glucose | - | 93 ± 0.1% |
| *Tecoma stans* (L.) Juss. ex Kunth | Bignoniaceae | 100 ± 0.00% |
| *Galeopsis segetum* Neck. | Lamiaceae | 97 ± 0.07% |
| *Tephrosia platycarpa* Guill. & Perr. | Fabaceae-Faboideae | 94 ± 0.09% |
| *Anogeissus leiocarpa* (DC.) Guill. & Perr. | Combretaceae | 93 ± 0.1% |
| *Cassia occidentalis* L. | Fabaceae-Caesalpinioideae | 93 ± 0.1% |
| *Ixora coccinea* L. | Rubiaceae | 90 ± 0.11% |
| *Volkameria inermis* L. | Lamiaceae | 90 ± 0.11% |
| *Duranta erecta* L. | Verbenaceae | 89 ± 0.11% |
| *Lantana camara* L. | Verbenaceae | 88 ± 0.12% |
| *Cassia alata* L. | Fabaceae-Caesalpinioideae | 84 ± 0.14% |
| *Albizia lebbeck* (L.) Benth. | Fabaceae-Mimosoideae | 77 ± 0.16% |
| *Azadirachta indica* A.Juss. | Meliaceae | 73 ± 0.18% |
| *Tridax procumbens* L. | Asteraceae | 73 ± 0.18% |
| *Antigonon leptopus* Hook. & Arn. | Polygonaceae | 70 ± 0.20% |
| *Jatropha integerrima* Jacq. | Euphorbiaceae | 68 ± 0.18% |
| *Sesbania pachycarpa* DC. | Fabaceae | 67 ± 0.21% |
| *Caesalpinia pulcherrima* (L.) Sw. | Fabaceae-Caesalpinioideae | 63 ± 0.22% |
| *Bignonia corymbosa* (Vent.) L.G.Lohmann | Bignoniaceae | 40 ± 28% |
| *Barleria prionitis* L. | Acanthaceae | 34 ± 29% |
| *Cassia sieberiana* DC. | Fabaceae-Caesalpinioideae | 29 ± 30% |
| *Urena lobata* L. | Malvaceae | 28 ± 29% |
| *Vernonia pauciflora* (Willd.) Less. | Asteraceae | 25 ± 28% |
| *Crotalaria retusa* L. | Fabaceae-Faboideae | 21 ± 30% |
| *Bougainvillea glabra* Choisy | Nyctaginaceae | 17 ± 33% |
| *Tradescantia pallida-purpurea* D.R.Hunt | Commelinaceae | 16 ± 32% |
| *Plumeria alba* L. | Apocynaceae | 14 ± 34% |
| *Senna corymbosa* (Lam.) H.S. Irwin & Barneby | Fabaceae-Caesalpinioideae | 14 ± 30% |
| *Cassia siamea* Lam. | Fabaceae-Caesalpinioideae | 10 ± 34% |
| *Canna indica* L. | Cannaceae | 9 ± 32% |
| *Combretum indicum* (L.) Jongkind | Combretaceae | 7 ± 35% |
| *Hyptis suaveolens* Poit. | Lamiaceae | 4 ± 37% |

**Supplementary Table 2: Mosquito** sample sizes used for each design, species, replicate, sex and diet (Experience 1.3)

| **Designs** | **Species (size)** | **Replicate (size)** | **Sex** | **5% glucose** | ***I. coccinea*** | ***C. pulcherrima*** | ***C. indicum*** |
| --- | --- | --- | --- | --- | --- | --- | --- |
| 1 | *An. coluzzii*  (687) | 1  (259) | M | 32 | 30 | 31 | 23 |
|  |  |  | F | 32 | 33 | 37 | 28 |
|  |  | 2  (247) | M | 34 | 32 | 33 | 31 |
|  |  |  | F | 30 | 27 | 32 | 28 |
|  |  | 3  (181) | M | 24 | 24 | 20 | 23 |
|  |  |  | F | 24 | 19 | 24 | 23 |
|  | *An. gambiae*  (766) | 1  (371) | M | 42 | 43 | 53 | 52 |
|  |  |  | F | 45 | 40 | 47 | 49 |
|  |  | 2  (223) | M | 28 | 28 | 25 | 26 |
|  |  |  | F | 32 | 28 | 26 | 30 |
|  |  | 3  (172) | M | 20 | 22 | 21 | 18 |
|  |  |  | F | 21 | 25 | 23 | 22 |
| 2 | *An. coluzzii*  (650) | 1  (281) | M | 33 | 38 | 35 | 35 |
|  |  |  | F | 34 | 32 | 30 | 44 |
|  |  | 2  (232) | M | 29 | 32 | 24 | 19 |
|  |  |  | F | 42 | 32 | 26 | 28 |
|  |  | 3  (137) | M | 18 | 12 | 17 | 08 |
|  |  |  | F | 23 | 17 | 22 | 20 |
|  | *An. gambiae*  (697) | 1  (320) | M | 27 | 31 | 37 | 45 |
|  |  |  | F | 53 | 57 | 35 | 35 |
|  |  | 2  (216) | M | 28 | 33 | 18 | 25 |
|  |  |  | F | 30 | 27 | 28 | 27 |
|  |  | 3  (161) | M | 19 | 24 | 18 | 15 |
|  |  |  | F | 23 | 21 | 22 | 19 |
| 3 | *An. coluzzii*  (690) | 1  (285) | M | 36 | 30 | 36 | 34 |
|  |  |  | F | 35 | 37 | 40 | 37 |
|  |  | 2  (248) | M | 47 | 37 | 30 | 33 |
|  |  |  | F | 34 | 26 | 18 | 23 |
|  |  | 3  (157) | M | 20 | 23 | 18 | 20 |
|  |  |  | F | 19 | 21 | 20 | 16 |
|  | *An. gambiae*  (674) | 1  (291) | M | 45 | 41 | 37 | 38 |
|  |  |  | F | 33 | 30 | 34 | 33 |
|  |  | 2  (213) | M | 25 | 27 | 32 | 26 |
|  |  |  | F | 26 | 34 | 17 | 26 |
|  |  | 3  (170) | M | 22 | 20 | 20 | 20 |
|  |  |  | F | 23 | 20 | 20 | 23 |

M = Males; F = Females

**Supplementary Table 3:** Statistical output of the multiple pairwise post-hoc comparisons (Experience 1.1)

| **Diet** | **Z. ratio** | **P-value** |
| --- | --- | --- |
| 5% glucose / *T. stans* | 0.02 | 1 |
| 5% glucose / *G. segetum* | 0.26 | 1 |
| 5% glucose / *T. platycarpa* | 0.31 | 1 |
| 5% glucose / *C. occidentalis* | 0.44 | 1 |
| 5% glucose / *A. leiocarpa* | 0.41 | 1 |
| 5% glucose / *V. inermis* | 0.84 | 1 |
| 5% glucose / *I. coccinea* | 0.42 | 1 |
| 5% glucose / *D. erecta* | 0.56 | 1 |
| 5% glucose / *L. camara* | 0.71 | 1 |
| 5% glucose / *C. alata* | 1.06 | 1 |
| 5% glucose / *A. lebbeck* | 1.65 | 1 |
| 5% glucose / *A. indica* | 2.37 | 0.88 |
| 5% glucose / *T. procumbens* | 2.42 | 0.86 |
| *5% glucose / A. leptopus* | 2.57 | 0.76 |
| 5% glucose / *J. integerrima* | 2.11 | 0.97 |
| 5% glucose / *S. pachycarpa* | 2.88 | 0.51 |
| 5% glucose / *C. pulcherrima* | 2.37 | 0.88 |
| 5% glucose / *B. corymbosa* | 4.16 | **0.01** |
| 5% glucose / *B. prionitis* | 4.44 | **0.004** |
| 5% glucose / *C. sieberiana* | 6.56 | **< 0.001** |
| 5% glucose / *U. lobata* | 5.30 | **< 0.001** |
| 5% glucose / *V. pauciflora* | 4.44 | **0.004** |
| 5% glucose / *C. retusa* | 6.82 | **< 0.001** |
| 5% glucose / *B. glabra* | 7.29 | **< 0.001** |
| 5% glucose / *T. pallida-purpurea* | 7.59 | **< 0.001** |
| 5% glucose / *P. alba* | 7 | **< 0.001** |
| 5% glucose / *S. corymbosa* | 7.03 | **< 0.001** |
| *5% glucose / C. siamea* | 7.18 | **< 0.001** |
| 5% glucose / *C. indica* | 7.65 | **< 0.001** |
| 5% glucose / *C. indicum* | 4.96 | **< 0.001** |
| 5% glucose / *H. suaveolens* | 6.12 | **< 0.001** |

**Supplementary Table 4:** Output of the statistical analyses of the effect of diet on the survival (without censoring) of mosquitoes (Experiment 1.2)

|  | **All species** | | | ***An. coluzzii*** | | | ***An. gambiae*** | | |
| --- | --- | --- | --- | --- | --- | --- | --- | --- | --- |
| Explanatory variables | LRT *X^2^* | df | P | LRT *X^2^* | df | P | LRT *X^2^* | df | P |
| Diet | 860.77 | 4 | **< 0.001** | 571.01 | 4 | **< 0.001** | 616.09 | 4 | **< 0.001** |
| Species | 2.52 | 1 | 0.11 | - | - | - | - | - | - |
| Sex | 10.60 | 1 | **0.001** | 12.12 | 1 | **< 0.001** | 2.17 | 1 | 0.14 |
| Diet: Species | 35.06 | 4 | **< 0.001** | - | - | - | - | - | - |
| Diet: Sex | 21.04 | 4 | **< 0.001** | 20.82 | 4 | **< 0.001** | 0.82 | 4 | 0.94 |
| Species: Sex | 4.04 | 1 | **0.04** | - | - | - | - | - | - |
| Diet: Species: Sex | 7.73 | 4 | 0.10 | - | - | - | - | - | - |

LRT = Likelihood Ratio Test; *χ^2^* = Chisq square; df = degree of freedom; P = P-value

**Supplementary Table 5:** Average longevity in days for all cages combined of *Anopheles coluzzii* and *Anopheles gambiae* males and females (±SE) maintained on one of 5 diet (Experiment 1.2, survival without censoring). Values represent Average longevity in days over the duration of the experiment until all mosquitoes are dead.

| **Species** | **Sex** | **Water** | **5% glucose** | ***I. coccinea*** | ***C. pulcherrima*** | ***C. indicum*** | **All diet** |
| --- | --- | --- | --- | --- | --- | --- | --- |
| *An. coluzzii* | Males | 2.29 ± 0.04 | 22.64 ± 1.10 | 8.89 ± 0.68 | 9.42 ± 0.54 | 9.42 ± 0.54 | 9.64 ± 0.41 |
|  | Females | 2.65 ± 0.05 | 21.35 ± 0.95 | 13.88 ± 0.91 | 11.30 ± 0.57 | 10.88 ± 0.54 | 12 ± 0.43 |
|  | Both sex | 2.46 ± 0.04 | 21.93 ± 0.72 | 11.13 ± 0.59 | 10.39 ± 0.40 | 9.01 ± 0.33 | 10.79 ± 0.30 |
| *An. gambiae* | Males | 2.43 ± 0.07 | 15.32 ± 1.20 | 9.89 ± 0.84 | 12.07 ± 0.65 | 6.34 ± 0.34 | 9.06 ± 0.40 |
|  | Females | 2.46 ± 0.05 | 18.01 ± 0.58 | 11.21 ± 0.66 | 13.13 ± 0.60 | 6.61 ± 0.46 | 10.05 ± 0.35 |
|  | Both sex | 2.45 ± 0.04 | 17.06 ± 0.57 | 10.69 ± 0.52 | 12.59 ± 0.44 | 6.48 ± 0.29 | 9.64 ± 0.26 |
| All species | Males | 2.34 ± 0.04 | 19.59 ± 0.87 | 9.28 ± 0.53 | 10.64 ± 0.43 | 7.09 ± 0.26 | 9.40 ± 0.29 |
|  | Females | 2.55 ± 0.04 | 19.62 ± 0.56 | 12.41 ± 0.56 | 12.09 ± 0.42 | 8.80 ± 0.40 | 11 ± 0.28 |
|  | Both sex | 2.46 ± 0.03 | 19.61 ± 0.48 | 10.92 ± 0.40 | 11.37 ± 0.30 | 7.90 ± 0.24 | - |

**Supplementary Table 6:** Statistical output of the multiple pairwise post-hoc comparisons (Experience 1.2, survival without censoring)

| **Diet** | **Z. ratio** | **P-value** |
| --- | --- | --- |
| Water / 5% glucose | 35.68 | **< 0.001** |
| Water / *I. coccinea* | 28.58 | **< 0.001** |
| Water / *C. pulcherrima* | 28.49 | **< 0.001** |
| Water / *C. indicum* | 24.18 | **< 0.001** |
| 5% glucose / *I. coccinea* | 13.49 | **< 0.001** |
| 5% glucose / *C. pulcherrima* | 14.12 | **< 0.001** |
| 5% glucose / *C. indicum* | 21.55 | **< 0.001** |
| *I. coccinea* / *C. pulcherrima* | 0.97 | 0.87 |
| *I. coccinea* / *C. indicum* | 9.58 | **< 0.001** |
| *C. pulcherrima* / *C. indicum* | 9.09 | **< 0.001** |

**Supplementary Table 7:** The effect of diet, species and sex on the proportion of mosquitoes tested positive to fructose (Experiment 1.2)

|  | **All species** | | | ***An. coluzzii*** | | | ***An. gambiae*** | | |
| --- | --- | --- | --- | --- | --- | --- | --- | --- | --- |
| Explanatory variables | LRT *X^2^* | df | P | LRT *X^2^* | df | P | LRT *X^2^* | df | P |
| Diet | 267.24 | 4 | **< 0.001** | 129.71 | 4 | **< 0.001** | 151.46 | 4 | **< 0.001** |
| Species | 2.78 | 1 | 0.1 | - | - | - | - | - | - |
| Sex | 2.96 | 1 | 0.09 | 2.19 | 1 | 0.14 | 0.87 | 1 | 0.35 |
| Diet: Species | 12.66 | 4 | 0.01 | - | - | - | - | - | - |
| Diet: Sex | 5.17 | 4 | 0.27 | 3.84 | 4 | 0.43 | 1.69 | 4 | 0.79 |
| Species: Sex | 0.92 | 1 | 0.34 | - | - | - | - | - | - |
| Diet: Species: Sex | 0.36 | 4 | 0.99 | - | - | - | - | - | - |

LRT = Likelihood Ratio Test; *χ^2^* = Chisq square; df = degree of freedom; P = P-value

**Supplementary Table 8:** The effect of diet, species, design, and their interactions on mosquito survivorship for all replicates (Experiment 1.3).

|  | **All designs** | | | **Design 1** | | | **Design 2** | | | **Design 3** | | |
| --- | --- | --- | --- | --- | --- | --- | --- | --- | --- | --- | --- | --- |
| Explanatory variables | LRT *X^2^* | df | P | LRT *X^2^* | df | P | LRT *X^2^* | df | P | LRT *X^2^* | df | P |
| Diet | 48.18 | 3 | **< 0.001** | 89.73 | 3 | **< 0.001** | 9.04 | 3 | **0.03** | 45.43 | 3 | **< 0.001** |
| Species | 1.75 | 1 | 0.19 | 0.88 | 1 | 0.35 | 6.82 | 1 | **0.01** | 0.01 | 1 | 0.92 |
| Sex | 0.12 | 1 | 0.72 | 0.46 | 1 | 0.50 | 1.60 | 1 | 0.21 | 5.78 | 1 | **0.02** |
| Design | 0.41 | 1 | 0.52 | - | - | - | - | - | - | - | - | - |
| Diet: Species | 6.44 | 3 | 0.09 | 6.83 | 3 | 0.08 | 4.31 | 3 | 0.23 | 4.81 | 3 | 0.19 |
| Diet: Sex | 4.60 | 3 | 0.20 | 1.83 | 3 | 0.61 | 13.21 | 3 | **0.004** | 12.40 | 3 | **0.01** |
| Species: Sex | 0.15 | 1 | 0.69 | 0.05 | 1 | 0.83 | 6.53 | 1 | **0.01** | 0.1 | 1 | 0.76 |
| Diet: Design | 0.15 | 3 | 0.001 | - | - | - | - | - | - | - | - | - |
| Species: Design | 0.29 | 1 | 0.59 | - | - | - | - | - | - | - | - | - |
| Sex: Design | 2.01 | 1 | 0.16 | - | - | - | - | - | - | - | - | - |
| Diet: Species: Sex | 1.32 | 3 | 0.72 | 0.39 | 3 | 0.94 | 3.73 | 3 | 0.29 | 5.63 | 3 | 0.13 |
| Diet: Species: Design | 6.53 | 3 | 0.09 | - | - | - | - | - | - | - | - | - |
| Diet: Sex: Design | 4.23 | 3 | 0.24 | - | - | - | - | - | - | - | - | - |
| Species: Sex: Design | 0.03 | 1 | 0.87 | - | - | - | - | - | - | - | - | - |
| Diet: Species: Sex: Design | 1.68 | 3 | 0.64 | - | - | - | - | - | - | - | - | - |

LRT = Likelihood Ratio Test; *χ^2^* = Chisq square; df = degree of freedom; P = P-value

**Supplementary Table 9:** Statistical output of the multiple pairwise post-hoc comparisons (Experience 1.3)

| **Diet** | **Z. ratio** | **P-value** |
| --- | --- | --- |
| 5% glucose / *I. coccinea* | 0.91 | 0.80 |
| 5% glucose / *C. pulcherrima* | 7.29 | **< 0.001** |
| 5% glucose / *C. indicum* | 13.86 | **< 0.001** |
| *I. coccinea* / *C. pulcherrima* | 6.36 | **< 0.001** |
| *I. coccinea* / *C. indicum* | 12.97 | **< 0.001** |
| *C. pulcherrima* / *C. indicum* | 7.04 | **< 0.001** |

**Supplementary Table 10:** Statistical analyses of the effect of diet, species, design, and their interactions on the insemination rate of mosquitoes for all replicates (Experiment 1.3)

|  | **All designs** | | | **Design 1** | | | **Design 2** | | | **Design 3** | | |
| --- | --- | --- | --- | --- | --- | --- | --- | --- | --- | --- | --- | --- |
| Explanatory variables | LRT *X^2^* | df | P | LRT *X^2^* | df | P | LRT *X^2^* | df | P | LRT *X^2^* | df | P |
| Diet | 25.38 | 3 | **< 0.001** | 23.14 | 3 | **< 0.001** | 41.14 | 3 | **< 0.001** | 33.91 | 3 | **< 0.001** |
| Species | 1.16 | 1 | 0.28 | 1.08 | 1 | 0.30 | 3.95 | 1 | 0.05 | 0.84 | 1 | 0.36 |
| Design | 3.97 | 2 | 0.14 | **-** | **-** | **-** | **-** | **-** | **-** | **-** | **-** | **-** |
| Diet: Species | 15.58 | 3 | **0.001** | 15.29 | 3 | **0.002** | 4.98 | 3 | 0.17 | 20.07 | 3 | **< 0.001** |
| Diet: Design | 16.08 | 6 | **0.01** | **-** | **-** | **-** | **-** | **-** | **-** | **-** | **-** | **-** |
| Species: Design | 3.14 | 2 | 0.21 | **-** | **-** | **-** | **-** | **-** | **-** | **-** | **-** | **-** |
| Diet: Species: Design | 14.64 | 6 | **0.02** | **-** | **-** | **-** | **-** | **-** | **-** | **-** | **-** | **-** |

LRT = Likelihood Ratio Test; *χ^2^* = Chisq square; df = degree of freedom; P = P-value

**Supplementary Table 11**: Insemination rate (±95CI) of mosquitoes for each diet treatment, species, and design across all replicates (Experiment 1.3)

| **Diet** | **All designs** | **Design 1** | **Design 2** | **Design 3** |
| --- | --- | --- | --- | --- |
| 5% glucose | 89 ± 0.03% | 82 ± 0.07% | 92 ± 0.04% | 93 ± 0.04% |
| *I. coccinea* | 80 ± 0.04% | 88 ± 0.06% | 87 ± 0.06% | 69 ± 0.09% |
| *C. pulcherrima* | 73 ± 0.05% | 76 ± 0.09% | 65 ± 0.11% | 77 ± 0.08% |
| *C. indicum* | 71 ± 0.06% | 78 ± 0.10% | 74 ± 0.12% | 67 ± 0.09% |
| All diet | - | 81 ± 0.04% | 82 ± 0.04% | 77 ± 0.04% |
| **Species** |  |  |  |  |
| *An. coluzzii* | 79 ± 0.03% | 81 ± 0.06% | 79 ± 0.06% | 78 ± 0.05% |
| *An. gambiae* | 80 ± 0.03% | 82 ± 0.05% | 85 ± 0.05% | 75 ± 0.06% |

**Supplementary Table 12:** Statistical output of the multiple pairwise post-hoc comparisons (Experience 1.3)

| **Diet** | **Z. ratio** | **P-value** |
| --- | --- | --- |
| 5% glucose / *I. coccinea* | 3.11 | **0.01** |
| 5% glucose / *C. pulcherrima* | 6.75 | **< 0.001** |
| 5% glucose / *C. indicum* | 7.11 | **< 0.001** |
| *I. coccinea* / *C. pulcherrima* | 3.80 | **< 0.001** |
| *I. coccinea* / *C. indicum* | 4.50 | **< 0.001** |
| *C. pulcherrima* / *C. indicum* | 1.15 | 0.66 |

**Supplementary Table 13:** Output of the statistical analyses of the effect of diet on the survival (with censoring) of mosquitoes (Experiment 1.2)

|  | **All species** | | | ***An. coluzzii*** | | | ***An. gambiae*** | | |
| --- | --- | --- | --- | --- | --- | --- | --- | --- | --- |
| Explanatory variables | LRT *X^2^* | df | P | LRT *X^2^* | df | P | LRT *X^2^* | df | P |
| Diet | 170.39 | 4 | **< 0.001** | 131.99 | 4 | **< 0.001** | 205.38 | 4 | **< 0.001** |
| Species | 0.36 | 1 | 0.55 | - | - | - | - | - | - |
| Sex | 0.58 | 1 | 0.45 | 0.44 | 1 | 0.51 | 0.92 | 1 | 0.34 |
| Diet: Species | 30.78 | 4 | **< 0.001** | - | - | - | - | - | - |
| Diet: Sex | 18.92 | 4 | **< 0.001** | 16.68 | 4 | 0.002 | 4.72 | 4 | 0.32 |
| Species: Sex | 1.43 | 1 | 0.23 | - | - | - | - | - | - |
| Diet: Species: Sex | 19.77 | 4 | **< 0.001** | - | - | - | - | - | - |

LRT = Likelihood Ratio Test; *χ^2^* = Chisq square; df = degree of freedom; P = P-value

**Supplementary Table 14**: Survival rate of *Anopheles coluzzii* and *Anopheles gambiae* males and females (±95CI) maintained on one of 5 diet (Experiment 1.2, survival with censoring). Values represent overall survival rates over the duration of the experiment until all mosquitoes are dead.

| **Species** | **Sex** | **Water** | **5% glucose** | ***I. coccinea*** | ***C. pulcherrima*** | ***C. indicum*** | **All diet** |
| --- | --- | --- | --- | --- | --- | --- | --- |
| *An. coluzzii* | Males | 10 ± 0.30% | 69 ± 0.20% | 43 ± 0.32% | 22 ± 0.33% | 6 ± 0.32% | 27 ± 0.14% |
|  | Females | 4 ± 0.27% | 92 ± 0.09% | 38 ± 0.29% | 41 ± 0.25% | 3 ± 0.31% | 33 ± 0.12% |
|  | Both sex | 7 ± 0.20% | 82 ± 0.10% | 40 ± 0.21% | 33 ± 0.20% | 4 ± 0.22% | 31 ± 0.09% |
| *An. gambiae* | Males | 5 ± 0.30% | 85 ± 0.11% | 46 ± 0.24% | 58 ± 0.22% | 27 ± 0.36% | 47 ± 0.11% |
|  | Females | 3 ± 0.25% | 81 ± 0.14% | 49 ± 0.22% | 41 ± 0.25% | 19 ± 0.31% | 35 ± 0.11% |
|  | Both sex | 4 ± 0.19% | 83 ± 0.09% | 47 ± 0.17% | 49 ± 0.17% | 22 ± 0.24% | 41 ± 0.08% |
| All species | Males | 8 ± 0.21% | 79 ± 0.10% | 45 ± 0.19% | 42 ± 0.19% | 14 ± 0.24% | 38 ± 0.09% |
|  | Females | 4 ± 0.18% | 87 ± 0.08% | 44 ± 0.18% | 41 ± 0.18% | 10 ± 0.22% | 34 ± 0.08% |
|  | Both sex | 5 ± 0.14% | 83 ± 0.07% | 44 ± 0.13% | 41 ± 0.13% | 12 ± 0.16% | - |

**Supplementary Table 15:** Statistical output of the multiple pairwise post-hoc comparisons (Experience 1.2, survival with censoring)

| **Diet** | **Z. ratio** | **P-value** |
| --- | --- | --- |
| Water / 5% glucose | 16.56 | **< 0.001** |
| Water / *I. coccinea* | 14.49 | **< 0.001** |
| Water / *C. pulcherrima* | 13.85 | **< 0.001** |
| Water / *C. indicum* | 3.48 | **0.005** |
| 5% glucose / *I. coccinea* | 8.10 | **< 0.001** |
| 5% glucose / *C. pulcherrima* | 8.92 | **< 0.001** |
| 5% glucose / *C. indicum* | 14.89 | **< 0.001** |
| *I. coccinea* / *C. pulcherrima* | 1.02 | 0.85 |
| *I. coccinea* / *C. indicum* | 11.55 | **< 0.001** |
| *C. pulcherrima* / *C. indicum* | 10.82 | **< 0.001** |

**Supplementary Table 16: Risk of mosquito survival (hazard ratio) with censoring for each treatment relative to the control (water).** lower .95 - upper .95 represents the 95% confidence interval around the hazard ratio (Experiment 1.2).

| **Diet** | **Hazard ratio (lower .95 - upper .95)** | **Z** | **P-value** |
| --- | --- | --- | --- |
| 5% glucose | 74.97 (0.01-0.02) | 16.82 | **< 0.001** |
| *Ixora coccinea* | 9.97 (0.07-0.14) | 14.29 | **< 0.001** |
| *Caesalpinia pulcherrima* | 9.05 (0.08-0.15) | 13.87 | **< 0.001** |
| *Combretum indicum* | 1.46 (0.54-0.87) | 3.12 | **0.002** |
